# Cities as parasitic amplifiers? Malaria prevalence and diversity in great tits along an urbanization gradient

**DOI:** 10.1101/2023.05.03.539263

**Authors:** Aude E. Caizergues, Benjamin Robira, Charles Perrier, Mélanie Jeanneau, Arnaud Berthomieu, Samuel Perret, Sylvain Gandon, Anne Charmantier

## Abstract

Urbanization is a worldwide phenomenon that modifies the environment. By affecting the reservoirs of pathogens and the body and immune conditions of hosts, urbanization alters the epidemiological dynamics and diversity of diseases. Cities could act as areas of pathogen dilution or amplification, depending on whether urban features have positive or negative effects on vectors and hosts. In this study, we focused on a host species and investigated the prevalence and diversity of avian malaria parasites (*Plasmodium/Haemoproteus* sp. and *Leucocytozoon* sp.) in great tits (*Parus major*) living across an urbanization gradient. In general, we observed high prevalence in adult birds (from 95% to 100%), yet lower prevalence in fledglings (from 0% to 38%). We found a slight tendency for increased Plasmodium sp. prevalence with increasing urbanization in adults. Urban nestlings had higher *Plasmodium* sp. infection rates than non-urban nestlings. We found evidence of higher diversity of parasites in the most natural urban park; however, parasite diversity was similar across other urbanization levels (e.g. from a little artificialized park to a highly anthropized industrial area). Parasite lineages were not habitat specific. Only one *Plasmodium* sp. lineage (YWT4) was associated with urban areas and some rare lineages (e.g., AFR065) were present only in a zoo area, perhaps because of the presence of African birds. This study suggests that urbanization can lead to a parasite amplification effect and can favor different avian malaria lineages.

## INTRODUCTION

Urbanization is a worldwide phenomenon driving environmental change and leading to the emergence of artificial habitats (Marzluff 2001; Gaston et al. 2015). Urban areas are a combination of remnant natural habitats and a complex assemblage of anthropogenic perturbations. They are characterized by new environmental conditions such as high levels of chemical, light, and sound pollution, increased impervious surfaces, and altered vegetation communities dominated by exotic plants (Forman and Godron 1986). Such extensive habitat modifications affect biodiversity at multiple ecological levels, from individual phenotypes to community assemblages. Notably, some species thrive in cities while others are not able to cope with urban conditions. Hence, urban communities are altered and mainly composed of fewer, often generalist and/or exotic, species with higher population densities compared to natural habitats (Shochat et al. 2006; Faeth et al. 2011).

Urbanization not only impacts individual species but also species interactions (Faeth et al. 2011), which can affect species evolution (Ots and Hõrak 1998; Marzal et al. 2005; Dyrcz et al. 2005). In particular, host-parasite interactions can be altered in urban habitats (Martin and Boruta 2013; Becker et al. 2015) because of variation in both the occurrence and abundance of species enabling the spread and transmission of the parasite (i.e., the vector species) (Reyes et al. 2013; Giraudeau et al. 2014; Neiderud 2015), changes in vectors’ feeding preferences in urban areas (Santiago-Alarcon et al. 2012; Abella-Medrano et al. 2018), and shifts in body condition and immune system efficiency of host species (Bailly et al. 2016; Capilla-Lasheras et al. 2017; Partecke et al. 2020). Depending on the positive and/or negative impact on the vector and host species, the effect of urbanization on disease prevalence can be twofold. First, in cases where urbanization predominantly negatively impacts vector species and/or predominantly favors the host species (e.g., if environmental requirements for parasite development are not met, Calegaro-Marques and Amato 2014), urban areas may act as a parasite dilution factor and urban animal populations should face lower risks of infections compared to their non-urban counterparts (Geue and Partecke 2008; Evans et al. 2009). Second, if the host species is predominantly negatively impacted by the urban conditions (e.g., immune depression in the host species, Bailly et al. 2016) urban individuals may suffer higher parasite burdens due to an amplification effect (e.g., Bichet et al. 2013).

Empirical evidence support both of these two scenarios, revealing case- and host-species dependence (Evans et al. 2009; Belo et al. 2011; Bichet et al. 2013b; Santiago-Alarcon et al. 2018). This might be because of the binary view of comparing urban *versus* non-urban habitats, with the postulate that the urban and non-urban environments stand as homogeneous and dichotomic environments. Yet, at a finer resolution, the urban matrix consists of a heterogeneous mosaic of local environments, some of which might be covered by impervious surfaces that contrast with green spaces. For example, parks offer great potential for multiple species to be supported (Nielsen et al. 2014; Lepczyk et al. 2017), sometimes leading to more diverse and species-rich areas than in nearby wild habitats (McKinney 2008). It therefore seems necessary to move from a binary perspective (i.e. the comparison between urban and non-urban habitats) to the study of a continuous urbanization gradient (e.g., French et al. 2008). Despite the growing body of literature on host-parasite interactions in urban habitats, their variations along an urbanization gradient are still poorly understood (Bradley and Altizer 2007; Delgado-V. and French 2012; Ferraguti et al. 2020).

Avian malaria parasites belong to *Haemoproteus*, *Plasmodium*, or *Leucocytozoon* genera and are widely studied in the context of host-parasite interactions (Rivero and Gandon 2018). Avian haemosporidians are ubiquitous parasites and encompass a vast diversity of species and strains that can be generalists, and infect a broad number of bird species (e.g. SGSI *Plasmodium relictum* strain), or specialist, and infect only one or few species (Valkiunas, 2004). They are vector-borne parasites infecting blood cells and mainly transmitted by five families of Diptera insects: *Culicidae*, *Hippoboscidae*, *Simuliidae*, *Ceratopogonidae*, and *Psychodidae* (Valkiunas and Iezhova 2018). These vectors are frequently encountered both in non-urban and urban areas, although their diversity and richness varies with habitat (Coene 1993). Indeed, the presence of water sources (river or pond) in urban areas is important for vector reproduction and population survival (Asghar et al. 2011). Among these vectors, some are known to be generalists and to feed on several vertebrate groups, especially in urban habitats (Jansen et al. 2009).

Great tits are common birds in Eurasia and are abundant in a wide range of habitats, from natural forests to heavily urbanized city centers (Fink et al. 2022). They are a good model species for ecologists and evolutionary biologists because they nest in human-provided nest boxes and are easy to capture and manipulate. Infection by avian malaria in Passeriformes is known to often induce an increase in immune response, lower survival, and reduced reproductive success (Ots and Hõrak 1998; Hõrak et al. 2001; Asghar et al. 2011; Lachish et al. 2011; Christe et al. 2012; Pigeault et al. 2018); therefore, if host-parasite interactions are affected by urbanization levels, the outcome for bird populations could depend on their habitat preference along the urban gradient.

In this study, we investigated the prevalence and diversity of avian malaria parasites in great tits (*Parus major*) in and around the city of Montpellier, south of France. Specifically, we (1) compared the prevalence in nestlings and adult individuals across different urbanization levels measured at the different scales, (2) characterized parasite molecular lineage richness and diversity along the gradient of urbanization, and (3) assessed the role of urbanization levels on parasite diversity.

## METHODS

### Study sites along an urbanization gradient

We studied nest boxes at two anthropogenically contrasted areas that had different levels of urban impacts. First the city of Montpellier, in southern France (43°36′N 3°53′E), which is a metropolitan area hosting 480,000 inhabitants. Second, the Rouvière oak forest, which is located 20 km northwest of Montpellier (Figure 1). In these city and forest contexts (hereafter urban and non-urban, respectively), long-term monitoring programs of the breeding populations of great tits have been conducted since 2011 and 1991, respectively (Charmantier et al. 2017). Monitoring consists of weekly visits mid-March to mid-July to document great tit reproduction in artificial nest boxes scattered in eight sites across the city (222 nest boxes) (Figure 1, see Figure S1 for a satellite image of the Zoo site) and across the forest of La Rouvière (94 nest boxes). The climate is typically Mediterranean, with mild winters and dry summers. Spring is marked by a sudden rise in temperature, coinciding with the great tit breeding season. This region of France hosts high densities of avian malaria *Plasmodium* vectors such as *Culex pipiens*, for which massive insecticide-based control treatments have been deployed for more than 60 years, with a focus on coastal areas (i.e., ca 15 km from Montpellier historical center, EID, 2020).

**Figure 1:**
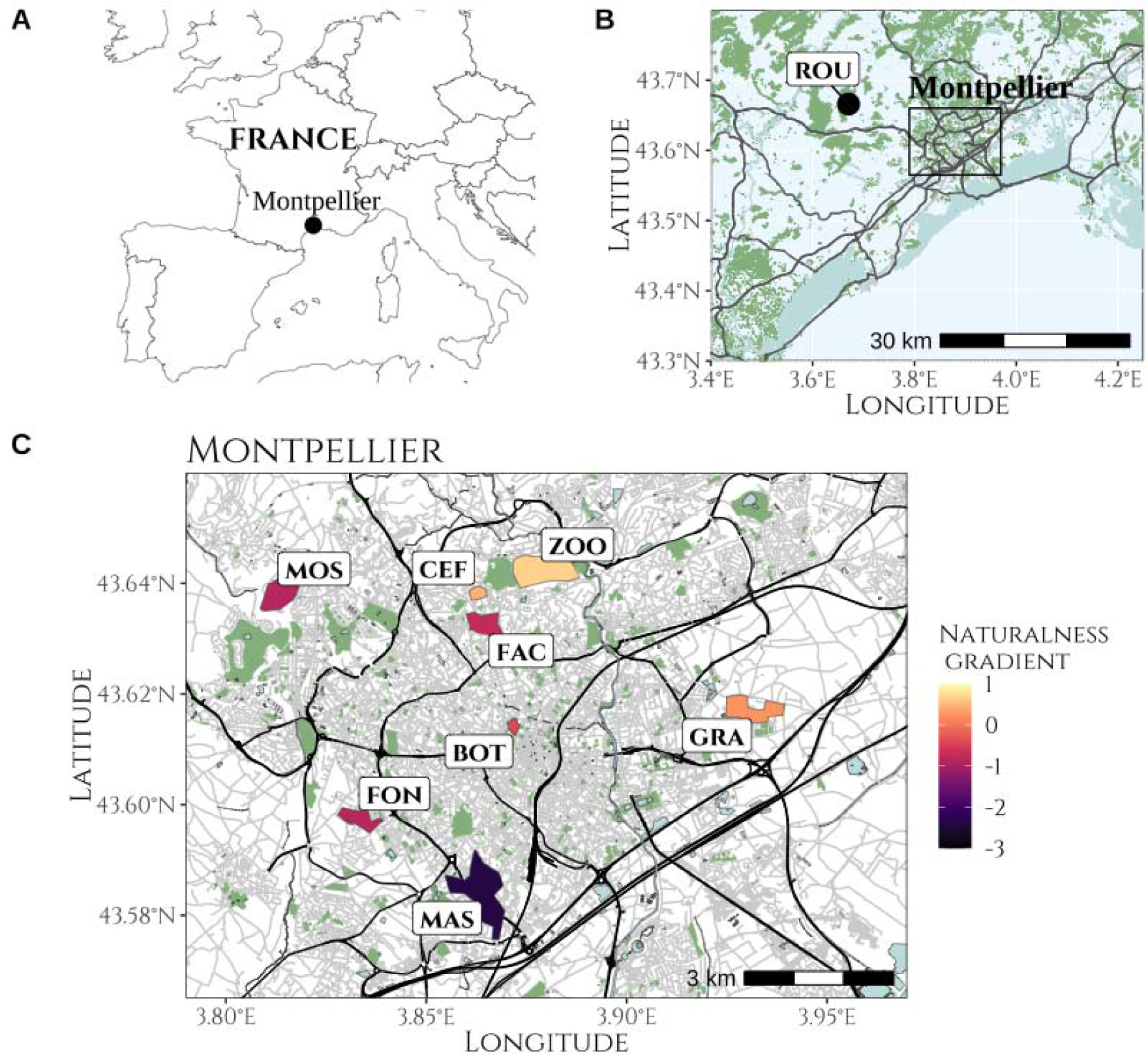
Maps of the sampling locations A) at European scale, B) at regional scale and C) at the city scale, where each polygon represents the limits of an urban sampling site and the colour represents the naturalness score of the site.

We characterized the level of urbanization and anthropogenic disturbance around each nest box, considering the area defined by a 50 m circular buffer around each nest-box where parents and nestlings were captured and sampled. This area is typically considered representative of a breeding great tit foraging area (Perrins 1979). We quantified four environmental features relevant for great tits breeding performances and fitness: (1) the extent of the vegetation cover (reflecting abundance of resources), (2) the motorized traffic disturbance (reflecting background noise pollution and chemical pollution), (3) the pedestrian disturbance (reflecting direct human disturbance), and (4) the amount of light pollution (affecting birds’ circadian rhythm, immunity and behavior). We measured the surface of vegetation cover (canopy and grass) around each nest box based on satellite images from Google maps. We quantified the motorized traffic perturbation by counting the number of motorized engines passing in the area during a 5 min count performed for each box in the early morning (between 7am and 11am). This count showed a 0.85 Pearson correlation with traffic data provided by the city of Montpellier (opendata.montpelliernumerique.fr/) in a given area (Demeyrier et al. 2016). We similarly estimated pedestrian disturbance with counts of pedestrians, bikes, and scooters. Finally, we defined local light pollution as the area covered by artificial light from lamp posts, assuming that a lamp post would illuminate a circular area of 50 m from its location. We summarized those four metrics using a principal component analysis as in Caizergues et al. (2021) to describe urbanization and disturbance at the nest level along two composite measures (Figure S2). In brief, we retained the two main axes explaining 67.8% of the variation in urban features, from which we used only the first axis in the present study. This first axis explained 42.4% of variance and was defined as the “naturalness” gradient, with positive values associated with larger vegetation cover, lower traffic disturbance, and lower light pollution. The second axis, defined as the “pedestrian frequency” gradient (25.4% of variance explained), was not used in the current study since it was not correlated with the habitat artificialization of an area but rather logically to the number of pedestrians passing by each nest box. We obtained site-level measures of “naturalness” for the eight urban sites and La Rouvière (sites hereafter referred to by acronyms made of their first three letters, see Table S1) by averaging these composite measures considering all nest boxes within a given site. This ranged from the most “natural” site, La Rouvière (ROU), to the most urbanized one, Mas Nouguier (MAS) in Montpellier city. Among urban sites, the zoo (ZOO; Figure 1C) was the least urbanized site, but was well settled in the urban matrix (Figure S1), with short distances from CEF and FAC allowing great tits from the zoo to interact with birds in these neighboring sites (see low genetic differentiation described in Perrier et al. 2017).

### Serologic sampling and molecular analyses

#### Blood sample collection

Between 2014 and 2019, we collected serologic samples between mid-March and mid-July. Samples were collected from 15 days old nestling and adult great tits across the urban and non-urban sites. We captured the parents when nestlings were 10-15 days old using traps inside nest boxes. All nestlings and adults were uniquely identified with rings provided by the Centre de Recherches sur la Biologie des Populations d’Oiseaux (CRBPO, Paris, France). We had a total of 296 adults (154 females 142 males) and 90 nestlings (not sexed and all sampled in 2014), see sampling detail Table S2.

We collected 10 µL of blood by performing a venipuncture in either the ulnar (i.e., wing) vein or a small subepidermal neck vein. We transferred blood samples using a capillary into an Eppendorf filled with 1 mL of Queen’s lysis buffer, then stored in 4°C refrigerators at the end of the field day until DNA extraction.

#### DNA extraction

We extracted total genomic DNA from blood samples using the DNeasy Blood and Tissue kit (Qiagen). We adapted the standard protocol by mixing 500 µL of solution of blood and Queen’s buffer (∼1/100 of blood) with 40 µL of proteinase K and 250 µL AL buffer. We then incubated the mixture at 56°C for 1.5 h. Afterwards, we added 8 µL of RNase A (100 mg/ml). We then performed DNA precipitation by adding 400 µL of ethanol.

#### Infection detection

We detected and identified parasites adapting Hellgren et al. (2004) protocol. We first amplified possible large fragments of mtDNA from *Plasmodium* spp., *Haemoproteus* spp. and *Leucocytozoon* spp. using polymerase chain reaction (PCR) with the HaemNF, HaemNR2 and HaemNR3 primers. PCR conditions included 15 min at 94°C, followed by 25 cycles of 30 s at 94°C, 40 s at 50°C, 1 min at 72°C, and a last cycle of 10 min at 60°C. Using 1 µL from the first amplified reaction, we then performed a secondary and more specific PCR to separately identify *Leucocytozoons* spp. and *Plasmodium*-*Haemoproteus* spp. presence with two different sets of primers: (i) we used the HaemF/ HaemR2 primers to amplify *Plasmodium* spp. And *Haemoproteus* spp. (test PH); (ii) and HaemFL/HaemRL primers to amplify *Leucocytozoon* spp. (test L). We performed this second PCR using Multiplex PCR kit Qiagen in a final volume of 10 µL following one cycle of 15 min at 94°C, 35 cycles of 30 s at 94°C, 40 s at 51°C/52°C (for *Leucocytozoon* spp.*/Plasmodium* spp. or *Haemoproteus* spp., respectively), 1 min at 72°C and one last cycle of 10 min at 60°C. We assessed amplification in 2% agarose gels leading to four possible infection outcomes: (1) uninfected (negative test PH and L), (2) infected by *Plasmodium* spp. and/or *Haemoproteus* spp. (positive test PH, negative test L), (3) infected by *Leucocytozoon* spp. (positive test L, negative test PH), and (4) coinfected by *Plasmodium* spp. and/or *Haemoproteus* spp. and *Leucocytozoon* spp. (positive test PH and L).

#### Lineage identification

We sent positive samples to Eurofins Genomics Company for Sanger sequencing. We then blasted sequences against the MalAvi database for molecular lineage identification (Bensch et al. 2009). We identified single and multiple infections of *Plasmodium* sp. and *Haemoproteus* sp. In contrast, the *Leucocytozoon* sp. sequencing quality was poor (i.e., there was an uncertain multiple-base identity in the sequence), and we were unable to identify a unique lineage (i.e., 100% blast score with a sequence from the database) for each sample. Therefore, we only identified a set of 5 likely lineages (blast >96%) for each sample. As no infection by any parasite from *Haemoproteus* genus was detected in our samples, we hereafter refer to *Plasmodium* sp. only.

### Statistical analyses

We performed all analyses with *R* software (version 4.2.1, R Core Team 2022). A complete list of the packages, associate versions, and reference used for data processing, analyses, and plotting, are provided in Supplementary Material Table S32.

#### Quantifying parasitic prevalence at the different sites

We estimated the site-level prevalence in nestlings and adults of *Plasmodium* sp. and *Leucocytozoon* sp. as the proportion of infected individuals as well as their 95% confidence intervals (Wilson score interval, “propCI*”* function of the *prevalence* package).

To further assess the role of urbanization in shaping prevalence patterns, we ran linear models separately on nestlings and adult individuals, and for the different parasite genera (*Plasmodium* sp. and *Leucocytozoon* sp., respectively), to link infection probability to the urban context across different spatial scales: the site level (i.e., average urbanization level of around all nests from a given site) and the local level (i.e., the urbanization level around the nest). To ease comparability with previous studies, we also carried out the analyses considering the habitat along the urban vs non-urban dichotomy (in this case the site and nest level always matched in their classification). To do so, we fitted three logistic regressions (“glm” and “glmer” function with a log-link function *stat* R package, Bates et al. 2015) with a binary response of infection (0 as not infected, 1 as infected) as a function of either habitat type (binary variable, 0 as non-urban, 1 as urban), the site-level naturalness (first axis of the PCA averaged on all the nest boxes of a sample site, see above), or the local nest-level naturalness (per nest-box first axis of the PCA value, and site as a random effect when the analysis was conducted considering nest-scale urbanization). For models ran on data from adult individuals, we further controlled for sex, age (in years), as well as the year of sampling. Because of the low number of samples in years 2017 (N = 6) and 2018 (N = 18), we removed these data from analysis to limit model convergence issues. We assessed the significance of each predictor using likelihood ratio tests (“drop1” function of the *stats* package) while dropping one predictor at a time.

We verified that linear models’ assumptions were not violated using various visual controls of residual distributions and associated statistical tests (histogram of residuals, Q-Q plot of expected residuals vs observed residuals, scattered plot of residuals vs estimates) using the *DHARMa* package as well as the *performance* package (see Supplementary Material: Supplementary text 1, Tables S2 to S31 and Figures S2 to S24). This raised no problem of collinearity, singular fit, convergence, or influential points.

#### Characterising lineage diversity and habitat specificity at the different sites

For subsequent analyses, we focused on adult individuals, as the quality of nestling malaria sequences was low and prevented us from correctly identifying lineages. Given the uncertainty in the *Leucocytozoon* sp. lineage identification (i.e., only a subset of likely lineages could be identified), we repeated the analyses (below) 1000 times for this parasite genus, each iteration randomly sampling a unique lineage (out of the 5 identified lineages) per individual. Thus, for *Leucocytozoon* sp. we provide the median estimates and associated 95% confidence intervals.

##### Lineage diversity

Haemosporidian lineages richness and abundance were analyzed with the *vegan* and *BiodiversityR* packages. To analyze patterns of lineage diversity per site, we estimated lineage richness and the combination between richness and evenness using the Shannon (roughly giving more weight to rare species) and inverse-Simpson (roughly giving more weight to common species) diversity indices. We also plotted the rank abundance curves for each study site, which highlight the richness (absolute value in the curve) and the evenness (slope of the curve) of parasite assemblages (Nagendra 2002).

We estimated dissimilarities in lineage composition between sites using the Bray-Curtis dissimilarity index (“vegdist” function of the *vegan* package). We computed this index on the binary sequence (i.e., indicating whether a given lineage was present or absent), and on the sequence of individual prevalence for each lineage (i.e., percent of infected individuals having the lineage). The former would provide insight into parasite composition resemblance (hereafter Bray-Curtis composition) and the latter into prevalence resemblance (hereafter Bray-Curtis prevalence).

##### Habitat specificity

To investigate whether some lineages occurred more frequently than randomly expected in urban versus non-urban environment, we compared the proportion of urban nest boxes at which each lineage was present to the overall number of nest boxes sampled using a binomial test (“binom.test*”* function). To avoid false negative habitat association in rare lineages, we computed the test only for lineages for which the type II error was below 0.20 (Cohen, 2013), and that occurred at least 10 times overall.

In addition, we investigated if parasitic community similarity was linked to urbanization at two scales: the sampling site and the nest box. We analyzed the correlation between naturalness distance (absolute difference in “naturalness” level) and parasite dissimilarity (Bray-Curtis composition) matrices using a mantel test with 999 permutations (“mantel.test” function of the *ape* package). We also controlled for spatial autocorrelation by testing whether parasitic community similarity was related to geographic proximity, repeating those analyses comparing the Euclidean distance between pairs of sites or nest boxes (“st_distance” function of the *sf* package) to parasite dissimilarity.

## RESULTS

### Plasmodium, Haemoproteus and Leucocytozoon prevalences

#### Parasitic prevalence in nestlings

In 15-day-old nestlings, avian haemosporidian prevalence was < 40% in both habitats, with some heterogeneity among sites (Figure 2A). No nestling was simultaneously infected by *Plasmodium* sp. and *Leucocytozoon* sp. parasites.

**Figure 2:**
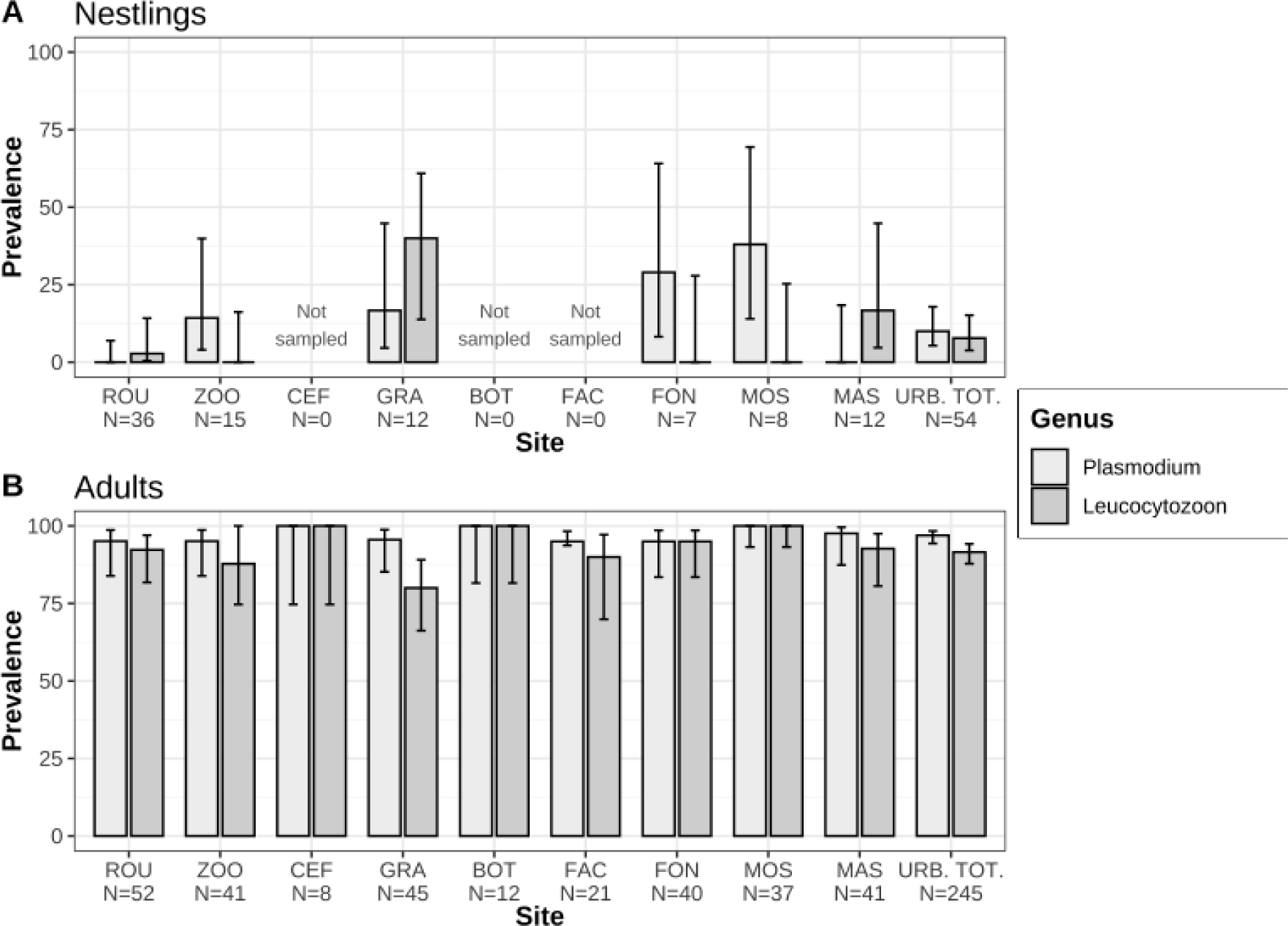
Mean prevalence of avian *Plasmodium* sp. (light grey) and *Leucocytozoon* sp. (dark grey) per site in great tit (A) nestlings and (B) adults. Error bars represent 95% confidence intervals. Sites are ordered by increasing urbanization level and sample size is detailed below each site.

The prevalence in *Plasmodium* parasites ranged from 0% to 38%, with an average of 16.33% (Figure 2A). Prevalence was significantly higher in the urban nestlings compared to non-urban nestlings (16.67% averaged on all urban sites vs. 0% in the non-urban site; *χ*^2^_1_ = 9.854, P = 0.002). However, overall small sample sizes and the presence of only one replicate of “non-urban” site potentially inflated these differences.). Plasmodium prevalence was not related to the nest- and site-level naturalness gradient (*χ*^2^_1_= 0.012, P = 0.908; *χ*^2^_1_= 1.186, P = 0.276, respectively).

The prevalence in *Leucocytozoon* sp. ranged from 0% to 40%, with an average of 9.90% and did not substantially differ consistently between urban and non-urban nestlings (11.11% averaged on all urban sites vs. 2.78% in the non-urban site; *χ*^2^_1_= 2.383, P = 0.123). *Leucocytozoon* sp. prevalence was unrelated to the nest- or site-level naturalness gradient (*χ*^2^_1_= 1.837, P = 0.175; *χ*^2^_1_= 1.291, P = 0.256, respectively).

#### Parasitic prevalence in breeding individuals

Avian haemosporidian prevalence ranged from 95% to 100% for *Plasmodium* sp. (mean = 97.04%), and 80% to 100% for *Leucocytozoon* sp. in breeding great tits, (mean = 92.93%) (Figure 2). Double infection was frequent (91.9% of individuals). In particular, all individuals infected with *Leucocytozoon* sp. were systematically infected with *Plasmodium* sp.

Prevalence of *Plasmodium* sp. and *Leucocytozoon* sp. did not vary significantly between urban and non-urban sites (*χ*^2^_1_= 0.003, P = 0.955; *χ*^2^_1_= 1.71, P = 0.191, respectively) nor with the site-level naturalness (*χ*^2^_1_= 0.360, P = 0.548; *χ*^2^_1_= 0.012, P = 0.911, respectively). However, nest-level naturalness gradient was potentially related to *Plasmodium* sp. prevalence (glmer: est. ± S.E. = -0.616 ± 0.397, *χ*^2^_1_= 2.937, P = 0.087), with a weak tendency for lower prevalence in less urbanized areas. In contrast, *Leucocytozoon* prevalence was not related to the nest-level naturalness gradient (*χ*^2^_1_= 0.567, P = 0.452). In addition, prevalence of both parasites genera did not vary between males and females (all P >> 0.05) or with age (all P >> 0.05). *Leucocytozoon* prevalence models showed a significant year effect when urbanization was considered dichotomous (glm: est. ± S.E. = 1.051 ± 0.484, *χ*^2^_1_ = 5.075, P = 0.024), with greater prevalence in 2019 compared to 2014. In contrast, *Plasmodium* sp. prevalence did not vary by year (all P >> 0.05).

#### Prevalence in nestlings versus breeding individuals

*Plasmodium* sp. and *Leucocytozoon* sp. prevalence at each site were not correlated between nestling and adult stages (Spearman correlation test, Il = 0.133, P = 0.803 and Il = -0.577, P = 0.231, respectively).

### Parasite molecular lineage diversity

A combination of 47 lineages of *Plasmodium* spp. and *Leucocytozoon* spp. species were recorded across all study sites (Figure 3), including 5 *Plasmodium* spp. and 42 *Leucocytozoon* spp. (total number of lineages identified by BLAST, not accounting for uncertainty in lineage identification). The *Plasmodium* sp. lineage SGS1 was the most represented of all lineages, with 272 infected birds out of 296 individuals sampled. Comparisons of lineage diversity depended on how diversity was quantified.

**Figure 3:**
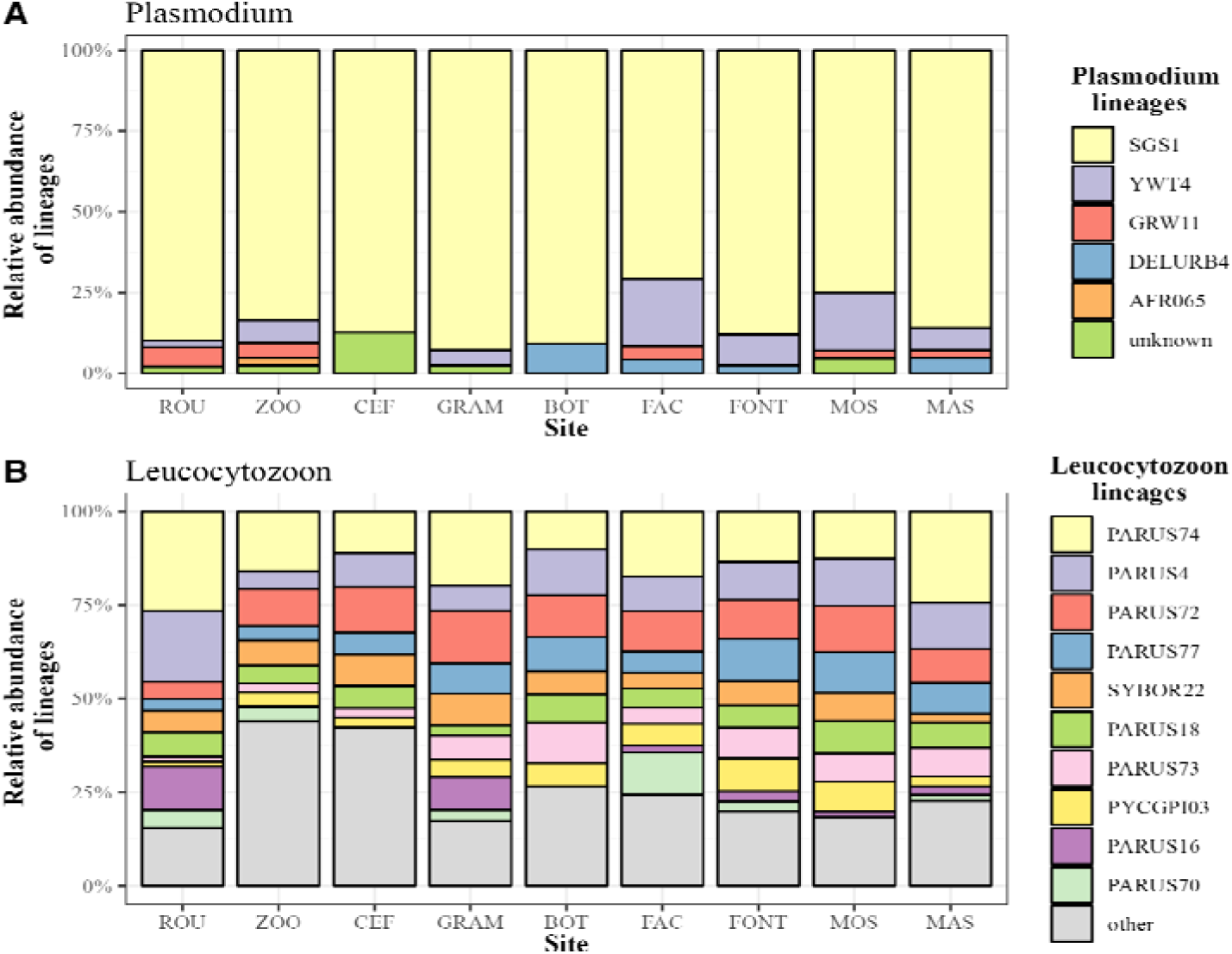
Proportions of (A) *Plasmodium* sp. and (B) *Leucocytozoon* sp. lineages found in each study site. For *Leucocytozoon* sp., only the most abundant lineages are shown in detail and lineages with less than 15 total occurrences were grouped as “other”.

The least urbanized urban site (ZOO) had the highest richness and Shannon’s evenness (richness = 22, evenness = 2.20, Table 1). The non-urban site (ROU) had intermediate richness (richness = 16) and was among the lowest in terms of Shannon’s evenness (evenness = 1.79). In contrast, FAC had the highest inverse Simpson’s diversity (Simpson’s index = 4.98). The non-urban site ROU had the lowest inverse Simpson’s diversity (Simpson’s index = 3.08, Table 1). Overall, diversity patterns among site were similar, with all sites having low evenness in parasite types which systematically consisted of a small subset of the 47 lineages (Figure 4).

**Figure 4:**
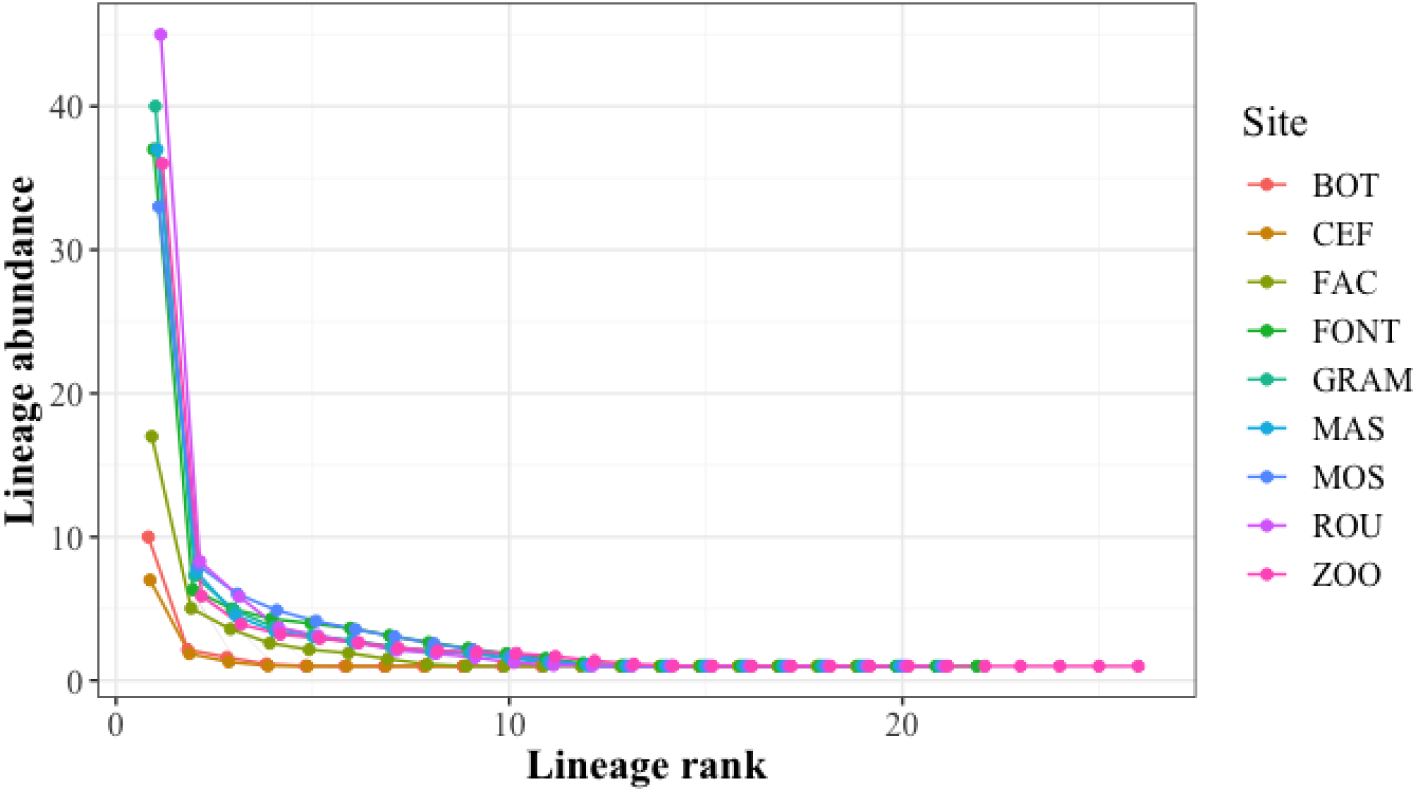
Rank-abundance curves for avian Haemosporidian lineages in each urban site and a non-urban site. Abundance is defined as the prevalence of a lineage at a given site. The *x*-axis represents the rank-abundance. The shape of the curve highlights the evenness: the steeper the curve, the less even distribution of lineage abundance. A flat curve indicates an evenly distributed community).

**Table 1:** Haemosporidian lineages richness and diversity indices (Shannon and Inverse Simpson) across the eight urban sites and the non-urban site (ROU).

| Site | N | Naturalness<br>index | Richness | Shannon | Inverse<br>Simpson |
| --- | --- | --- | --- | --- | --- |
| MAS | 41 | -2.383 | 18<br>(15 - 20) | 1.98<br>(1.88 - 2.06) | 3.65<br>(3.56 - 3.72) |
| MOS | 37 | -0.865 | 16<br>(14 - 19) | 2.09<br>(1.99- 2.17) | 4.53<br>(4.41 - 4.61) |
| FON | 40 | -0.854 | 18<br>(16 - 21) | 2.09<br>(2.00 - 2.17) | 4.11<br>(4.02 - 4.17) |
| FAC | 21 | -0.750 | 15<br>(13 - 17) | 2.14<br>(2.01 - 2.23) | 4.98<br>(4.77 - 5.13) |
| BOT | 12 | -0.406 | 9<br>(7 - 11) | 1.71<br>(1.54 - 1.84) | 3.51<br>(3.33-3.64) |
| GRA | 45 | 0.254 | 16<br>(13 - 18) | 1.86<br>(1.76 - 1.94) | 3.33<br>(3.25 - 3.38) |
| CEF | 8 | 0.458 | 8<br>(6 - 9) | 1.71<br>(1.49 - 1.80) | 3.81<br>(3.46 - 3.95) |
| ZOO | 41 | 0.687 | 22<br>(19 - 25) | 2.20<br>(2.10 - 2.28) | 4.11<br>(4 - 4.17) |
| ROU | 52 | 1.221 | 16<br>(14 - 19) | 1.79<br>(1.69 - 1.88) | 3.08<br>(3.02 - 3.13) |

### Habitat specificity of lineages in breeders

Regarding lineage habitat specificity, we found one lineage, YWT4 (*Plasmodium* sp.), that occurred more in urban habitats than expected by chance (Figure 5). None of the other *Plasmodium* sp. or *Leucocytozoon* sp. lineages were statistically more associated with one habitat type than the other.

**Figure 5:**
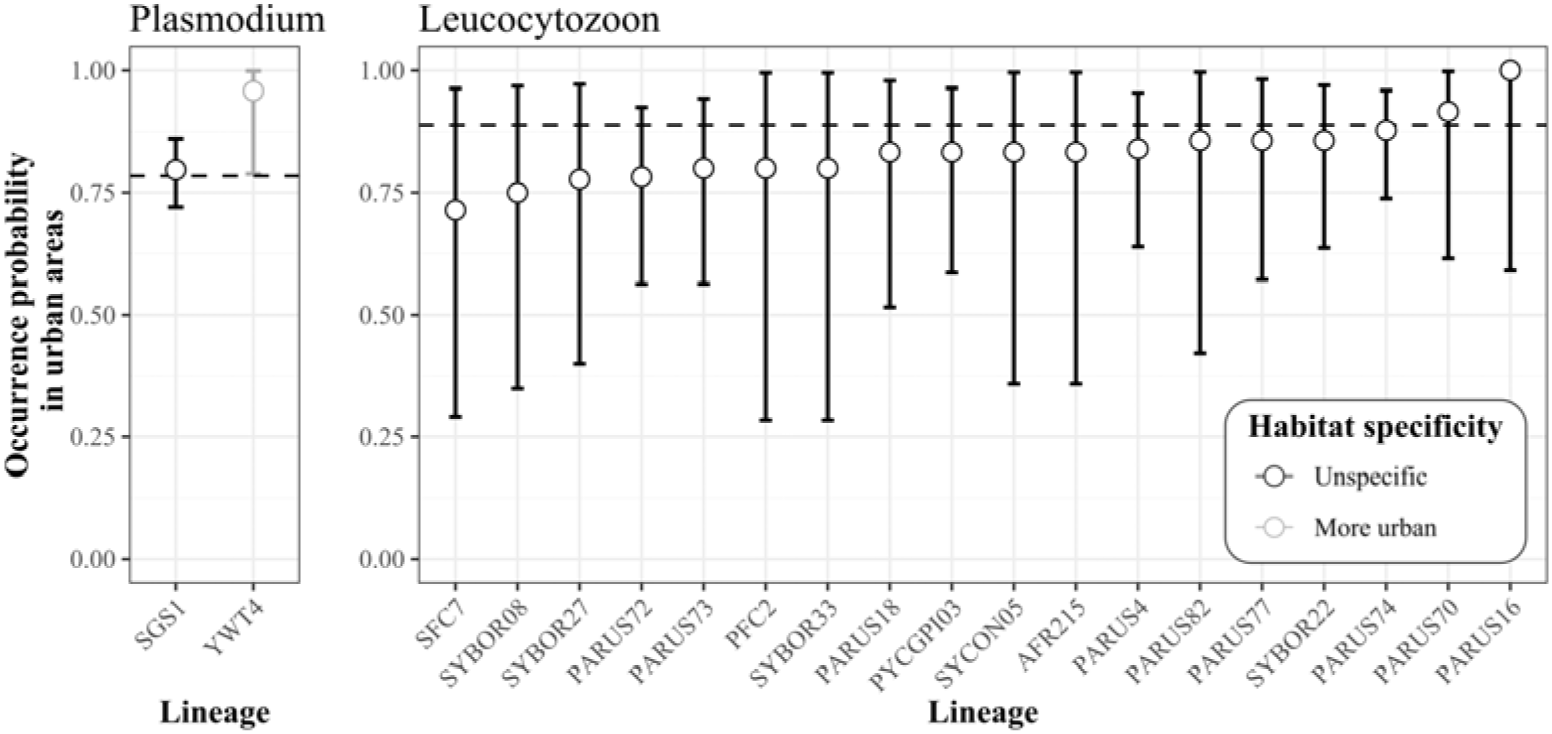
Occurrence probability of avian Haemosporidian lineages in the urban habitat for A) *Plasmodium* sp. and B) *Leucocytozoon sp*. Error bars represent 95% confidence intervals. The dashed line represents the expected probability of occurrence of a lineage in the urban habitat under random distribution. Grey dots and error bars represent lineages that are found statistically more in the urban habitat, and black, lineages that are not habitat-specific.

Resemblances between sites were globally homogenous between pairs of sites (Figure 6), both in composition (i.e., in terms of lineage diversity) and prevalence (i.e., in terms of infection rate for a given lineage). Anecdotically, BOT and CEF, the smallest and least sampled sites, were the most dissimilar to other sites (Figure 6).

**Figure 6:**
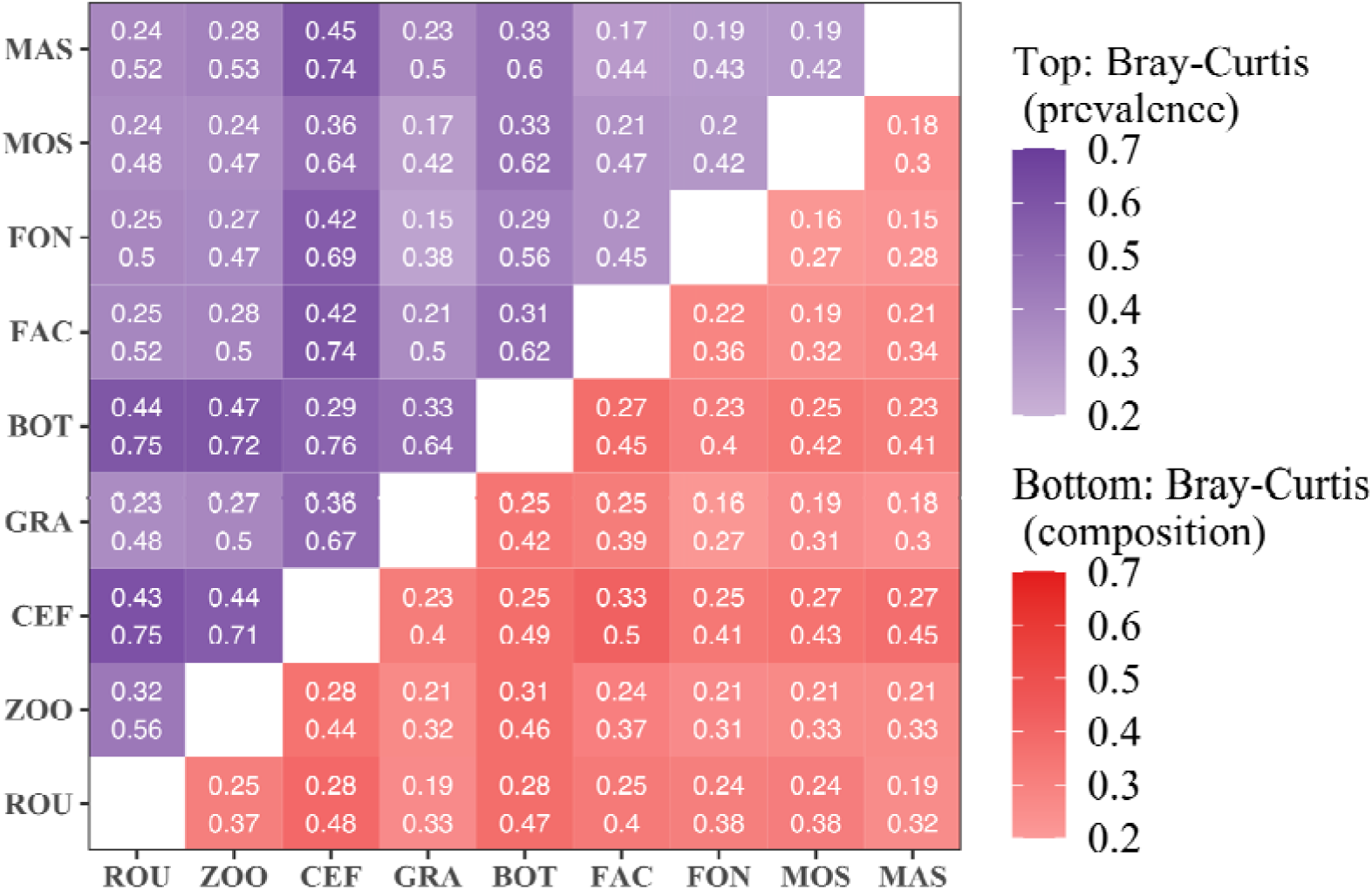
Heatmap of Bray-Curtis dissimilarity between each site considering binary sequences of lineage composition (bottom) or the prevalence of each lineage among infected individuals (top). Darker colors represent higher values of Bray Curtis index and stronger differences in lineage composition/prevalence between a pair of sites. Values in the cells indicate the upper border of the 95% confidence interval.

We found no statistical link between parasitic community similarity and naturalness gradient or geographical proximity at both the site or the nest box levels (Mantel test: P >> 0.05 for all the 1000 subsampled datasets; p-values were adjusted to maintain the false discovery rate to 5%).

## DISCUSSION

In this study, we investigated the link between urbanization and avian malaria prevalence and lineage diversity at different scales across wild populations of great tits in and around Montpellier, a metropolis of almost half a million inhabitants in southern France. We found marked differences in parasite prevalence between life stages, with 15-day-old nestlings showing substantially lower parasite prevalence than adult birds. Malaria parasite prevalence also varied depending on the environment, with urban nestlings significantly more infected than non-urban nestlings. There was also a tendency for a higher *Plasmodium* prevalence in adults from more urbanized nests. Altogether, this suggests the possible parasitic amplification effect in the city. However, the overall parasite diversity was unvariant across urbanization contexts. Only some haemosporidian lineages occurred solely or more often in urban areas, suggesting the possibility for habitat specificity of parasite strains.

### Life stage and habitat-dependent prevalence

Overall, infection by *Plasmodium* was greater than infection by *Leucocytozoon*, which is a common pattern observed across bird species (Pigeault et al. 2018, but see Merino et al. 2008). Haemosporidian prevalence was overall low in nestlings (from 0% to 38%) but high in adults (from 95% to 100%), and this pattern was consistent in both urban and non-urban areas. Such prevalence levels are comparable to previous studies for adult great tits (Glaizot et al. 2012; Rooyen et al. 2013). To our knowledge, this is the first time it is tested in 15 day-old urban great tit nestlings. In fact, only lower prevalence in young juvenile (one year-old) birds compared to adults was previously described in great tits and other passerine species (Wood et al. 2007; Santiago-Alarcon et al. 2016). The higher infection detection in adults than in nestlings frames coherently with the vector (e.g., *Culex pipiens*) life cycle, with a progressive increase in adult mosquitos and associated infection risk from spring to summer (Zélé et al. 2014). As a consequence, the risk for 15 day-old nestlings of being infected is expected to be low as they were sampled during spring. Similarly, Valkiunas and Iezhova (2018) found that young adults presented lower prevalence, which is in line with the fact that Haemosporidian infections yield an acute infection followed by a life-long chronic infection. Hence, the longer the exposure to the parasites, the higher the probability of eventually being infected. Possibly, the lower infection in 15 day-old nestlings could also be due to the delay of detection that is not immediate after infection (Cosgrove et al. 2006).

Our results tend to support the hypothesis of the existence of a parasitic burden in more urbanized areas. Specifically, the observed statistical effect of nest-level urbanization on *Plasmodium was* marginal. It would thus deserve further attention to clearly determine the role of the urbanization at such scale. This potential effect would contrast frequent reports of lower parasitic prevalence in urban areas including in our focal species (Bailly et al. 2016). Whether avian malaria is more or less prevalent in cities thus appears strongly case-specific (Evans et al. 2009).. Differences in parasite prevalence between habitats may be directly induced by variations in the presence and/or density of vectors (e.g., Martínez-de la Puente et al. 2013). These variations should be the consequence of presence or absence of their suitable ecological niches. For instance, among the 11 paired populations of blackbirds *Turdus merula* studied by Evans et al. (2009), in 3 cases, avian malaria prevalence was found to be higher in urban areas as a consequence of underwater area presence. While fine scale densities of vectors are not yet known for the city of Montpellier and its surrounding area, a tendency towards higher malaria prevalence in more urbanized areas could indicate higher population size or densities of vectors in such areas, perhaps given the marshes nearby. This, however, remains to be empirically demonstrated.

In addition, urban nestlings showed higher prevalence than non-urban ones, although the non-urban site had only one replicate, limiting robust generalization. Replication of the work done here at other sites is therefore crucial to (in)validate this finding. Provided that this is generalizable, one can hypothesize that early malaria infection in urban nestlings might be an indirect result of the heat island effect. Indeed, Paz and Albersheim (2008) showed that higher temperatures in urban areas proved beneficial to *Culex pipiens* mosquitoes growth and that some diseases (i.e., the human West Nile Fever) transmitted by this vector appeared earlier in the season in the city compared with surrounding countryside areas. Hence, environmental shifts observed in urban areas can be directly linked to spatial and temporal parasite infections. In addition, malaria infections are known to vary in time (Zélé et al. 2014). Given the role of the urban area in buffering on climatic variations, urbanization could be responsible for major changes in seasonality of parasitic infection. As shown here, this could cascade onto the emergence of earlier disease outbreak and earlier nestling contamination. The link between urban specific climatic features and seasonality of vectors and disease outbreaks in urban areas remains overlooked and should be the focus of further research avenues.

### Spatial heterogeneity in lineage diversity

When exploring diversity of Haemosporidian lineages across sites, we mostly found similar levels of diversity along the urbanization gradient and no strong ‘cluster’ of similar lineages in similarly urbanized or closer sites. Anecdotically, the non-urban sites had the lowest Haemosporidian lineage diversity, whereas the large zoo urban park had the highest. Interestingly, previous studies reported that urban parks with higher diversity of plant and bird species were also the most diverse in terms of Haemosporidian lineages (in multiple species: Carbó-Ramírez et al. 2017; in the House Sparrow: Jiménez-Peñuela et al. 2021). In our case, the Zoo du Lunaret consists of an 80-ha natural area where a large diversity of both native and exotic plant and bird species coexist. Interestingly, the only occurrence of *Plasmodium* sp. AFR065 lineage was in this zoo. According to the *MalAvi* database (Bensch et al. 2009), this lineage was found previously only on the African continent, in two bird genus in Malawi (*Cercotrichas* and *Andropadus,* Lutz et al. 2015). Hence, the presence of such lineages in this particular area of the city is most probably linked to the presence of captive African birds in the zoo (see next section for details on these birds).

The diversity of Haemosporidian lineages at the non-urban site of La Rouvière, 20 km away from the city of Montpellier, ranked among the lowest in richness and evenness (Table 1 and Figures 3 and 4), which contrasts with previous results found and showed opposite trends when comparing urban and non-urban sites (e.g., in the House Sparrow : Jiménez-Peñuela et al. 2021). The difference in diversity highlighted by these indices may however be biologically small, as the dissimilarity between ROU and the other sites was in the range of any other pairs of sites. In our study site, the overall urban habitat presents numerous ornamental plant species, whereas the non-urban habitat, which is a Mediterranean forest, is mainly dominated by oak trees. Hence, even with lower density of vegetation, the urban areas might be prone to a maintain high diversity of pathogens (Carbó-Ramírez et al. 2017). However, such hypothesis remains to be further tested. In particular, since vector distribution, abundance, and diversity are likely to play a major role in malaria infection patterns observed in bird hosts, disentangling the processes underpinning parasite prevalence and diversity patterns in different urban conditions will require combining parallel investigation of vectors and hosts along gradients of urbanization. In this study, we however only focused on the host.

### Habitat specific lineages

While none of the sampled sites revealed a striking divergent composition in Haemosporidian lineages, hence similar diversity trends, we still observed some heterogeneity in lineage type occurrence. Overall, the *Plasmodium* sp. infections were mainly dominated by SGS1 lineage (*Plasmodium relictum*). SGS1 is known to be a generalist lineage, present in multiple avian species and environments (Rooyen et al. 2013) and transmitted by *Culex pipiens* (Ventim et al. 2012; Inci et al. 2012), which is widely present in the south of France in both habitats. Aside from SGS1, some lineages were found in low occurrence exclusively in the urban habitat: AFR065 occurred once in ZOO and DELURB4 occurred in urban sites only. Habitat specificity analyses controlling for unequal sampling across the sites revealed that only one *Plasmodium* sp. lineage (YWT4) occurred more in urban habitats. No lineage was found to be associated with the forest habitat. Yet, this is possible that it is due to a lack of power in our forest dataset as it included only NN individuals. When investigating the previous occurrences of the 3 specific urban associated lineages (i.e., AFR065, DELURB4 and YWT4) in the MalAvi database, we found that they were relatively rarely encountered, at least in great tits.

AFR065 was reported only twice, once in the Miombo scrub robin (*Cercotrichas barbata*, Muscicapidae) and once in the western greenbul (*Andropadus tephrolaemus,* Pycnonotidae) in Malawi and never on the European continent nor in the great tits (Lutz et al. 2015). As mentioned before, the individual infected by AFR065 was captured in the Zoo du Lunaret (most natural urban site). At the time of the sampling for this study, the zoo hosted 65 African birds from 14 different species. While malaria infection status of these captive birds held in the zoo are low (<5%, unpublished data), we can hypothesize that they were the initial carriers of AFR065 that was then transferred to a great tit via the contaminated vectors. This result raises concern regarding local wildlife epidemiology when introducing or keeping exotic wildlife captive in contact with native species.

We found no previous occurrence of the DELURB4 lineage in great tits in the *MalAvi* database, even if this lineage was previously shown to be the second most common lineage present in the vector *C. pipiens* in the area (Zélé et al. 2014), and numerously recorded in the close sister species the Blue tit *Cyanistes caeruleus* (Ferrer et al. 2012) and in other bird families (e.g., *Passeridae*, *Turdidae* and *Muscicapidae*) in several European countries (Spain, Italy, Bulgaria, Russia according to the *MalAvi* database). Similarly, YWT4 is a rare lineage with only 7 occurrences in the whole *MalAvi* database, mainly in the Western yellow wagtail (*Motacilla flava*), but was found 25 times in the studied urban great tits, and once in a non-urban bird. Reasons why these lineages were more common in urban areas than in non-urban habitats remain to be explored. A possible explanation could be the difference in bird community composition between habitats, leading to contact with different bird species, each with their own body of specific Haemosporidian parasite lineages as suggested by the occurrence of a rare lineage in the zoo that is hosting African species. Testing this hypothesis would require a thorough scan of Haemosporidian infections in multiple host species from both urban and non-urban habitats in replicated cities combined with a thorough characterization of bird community assemblages and abundances along urbanization gradients.

## Conclusion

While we found no striking difference in malaria prevalence between urban and non-urban great tits, urbanization was associated with earlier infections in nestlings. In addition, we found a weak tendency for *Plasmodium* sp. prevalence to increase with urbanization. While our results will need to be replicated with higher number of sampled sites and individuals, they could suggest that urbanization does not decrease parasitic load but may, on the contrary, lead to a parasitic burden for urban great tits. Interestingly, although sites displayed no major differences in Haemosporidian lineage community composition, urban sites hosted preferentially lineages that rarely occurred in malaria databases. This suggests that urbanization could play a role in the emergence and spread of previously rare disease strains, especially when zoos are present.

## STATEMENTS AND DECLARATIONS

### Ethical statement

Captures were performed under personal ringing permits delivered by the CRBPO (Centre de Recherches par le Baguage des Populations d’Oiseaux, e.g., ringing permit for Anne Charmantier number 1907) for the Research Ringing Programme number 369. All experimental protocols were approved by the ethics committee for animal experimentation of Languedoc Roussillon (CEEA-LR, most recent approval in 2018 for APAFIS#8608-2017012011062214) as well as by Regional Institutions (most recent bylaw issued on 07/04/2022 by the Prefecture n° 2B-2022-04-07-00002).

### Data and code sharing

Data and code used for this study are freely available on Zenodo via Github (DOI : 10.5281/zenodo.8329693 & https://github.com/AudeCaizergues/Malaria_Great_Tits).

### Authors contribution

A.E.C., S.P & A.C. collected the samples along with field collaborators. M.J. & A.B. performed the molecular analyses. A.E.C. & B.R. conducted the statistical analyses and wrote the manuscript. C.P., S.G. & A.C. conceptualised the research. S.G. & A.C. financed the project. All authors contributed to writing the manuscript.

## Acknowledgements

We are grateful to the managers and the employees of the Zoo de Lunaret, Montpellier, especially Baptiste Genet, David Gomis, Marc Romans as well as the PLT platform of the CEFE for their help in data collection and their feedback on our research. We also thank the city Council of Montpellier for permitting us to carry out this long-term research project.

