## Supplementary materials for "Cities as parasitic amplifiers? Malaria prevalence and diversity in great tits along an urbanization gradient"

### 1.  Site Locations and sample sizes

**Table S1:** Correspondence of acronyms, full name and location of studied sites.

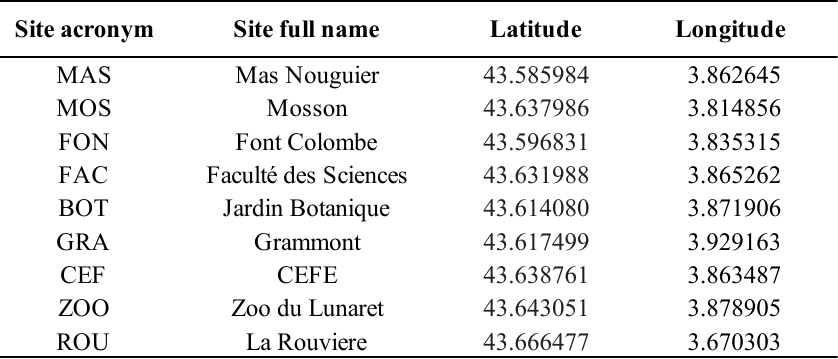

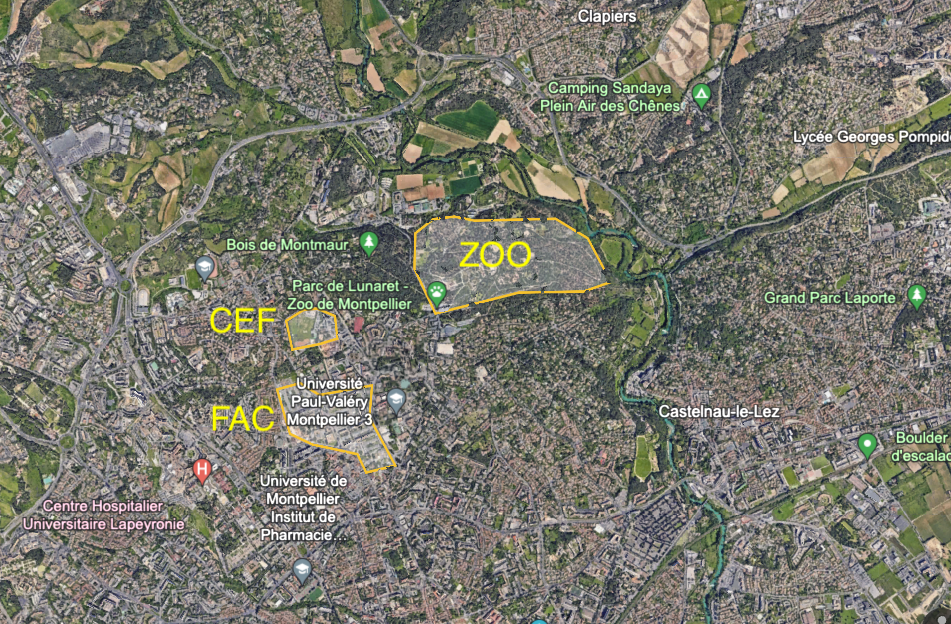

**Figure S1:** Satellite view of the Zoo area | The image was taken from Google Earth Engine (Gorelick et al. 2017) on the 20/11/2023. The Zoo site is located close to urban sites (e.g., about 600 m away from the CEF site; average distance between all urban sites is 4.4 km). In comparison, the forest site is located 20 km away from the city, in a rural landscape.

**Table S2:** Sample size per site and year in nestlings and adults, and associated sex ratio (in adults only).

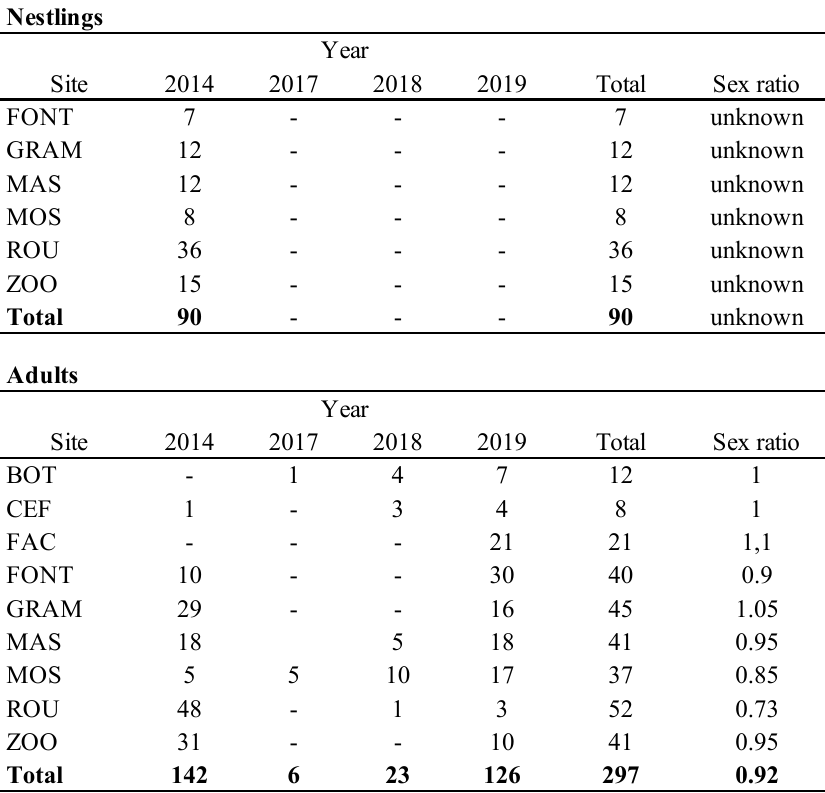

### 2. PCA on environmental factors

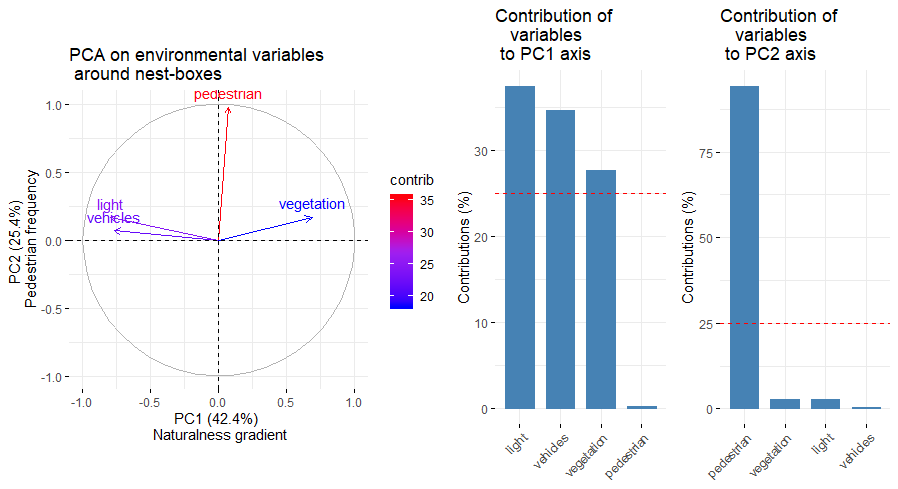

A

B

**Figure S2:** (A) Results from the PCA analysis on environmental factors measured around nest-boxes: vegetation cover, vehicle density, pedestrian frequency and light pollution. (B) Histograms showing the contribution of each variable to the first two Principal Component axes. These histograms show that PC1 is interpretable as a naturalness gradient and PC2 as pedestrian disturbance (from Caizergues et al. 2021).

### 3. Model testing functions

**Supplementary text 1:** Functions used to perform diagnostics on the mixed models. For each model we present an overview of the model results as well as the verification of the model assumption and stability. Overall, we observed high stability despite some small deviations of assumptions. In particular, histograms of residuals were skewed, a likely consequence of the small sample size, but this was inconsequential (see associated QQ-splot and further stability assessment). Such deviations are thus unlikely to have affected our estimations (Schielzeth et al., 2020).

A) **Function to check linear model assumptions**
   a. **Collinearity plot**

```
as_huxtable(model_name$collinearityTest %>% as.data.frame()) %>%
  set_caption("Collinearity assessment of the model ") %>%
set_all_padding(4) %>%
set_outer_padding(0) %>%
set_number_format(2) %>%
set_bold(row = 1, col = everywhere) %>%
set_bold(col = 1, row = everywhere) %>%
set_bottom_border(row = 1, col = everywhere) %>%
set_width(1.2) %>%
as_flextable()
```

b. **Distribution of residuals**

```
diagnostics.plot.dharma <-
  function(mod.res,
           col = grey(level = 0.25, alpha = 0.5),
           breaks.histo = 20,
           quantreg = TRUE) {
    old.par = par(no.readonly = TRUE)
    par(mfrow = c(2, 2))
    par(mar = c(3, 3, 3, 0.5))
    hist(
      residuals(mod.res),
      probability = T,
      xlab = "",
      ylab = "",
      main = "",
      breaks = breaks.histo )
    mtext(text = "Histogram of residuals",
          side = 3,
          line = 0)
    x = seq(min(residuals(mod.res)), max(residuals(mod.res)), length.out =
              100)
    lines(x, dnorm(x, mean = 0, sd = sd(residuals(mod.res))))

    library(DHARMa)
    simulationOutput <- simulateResiduals(fittedModel = mod.res, plot = FALSE)
   	plotQQunif(simulationOutput) # left plot in plot.DHARMa()
    	plotResiduals(simulationOutput, quantreg = quantreg)
    	par(old.par)
  }
```

B) **Stability plot**

as_huxtable(model_name$dfbetas %>% as.data.frame()) %>%

set_caption("Dfbetas of the model ") %>%

set_all_padding(4) %>%

set_outer_padding(0) %>%

set_number_format(2) %>%

set_bold(row = 1, col = everywhere) %>%

set_bottom_border(row = 1, col = everywhere) %>%

set_width(1.2) %>%

as_flextable()

### 4. Nestling models

#### 4.1. Urbanization as binary variable (urban vs. rural)

##### 4.1.1. Linear model results

**Table S3:** Detailed output of fixed effects included in the GLM investigating *Plasmodium/Haemoproteus* prevalence in nestlings with urbanization as a binary variable.

|  | **Infection (Plasmodium/Haemoproteus)** | | | |
| --- | --- | --- | --- | --- |
| *Predictors* | *Log-Odds* | *std. Error* | *CI* | *p* |
| (Intercept) | -19.57 | 1792.34 | -572.61 – 63.86 | - |
| Habitat [Urban] | 17.96 | 1792.34 | -131.38 – NA | 0.002 |
| Observations | 90 | | | |

**Table S4:** Detailed output of fixed effects included in the GLM investigating *Leucocytozoon* prevalence in nestlings with urbanization as a binary variable.

|  | **Infection (Leucocytozoon)** | | | |
| --- | --- | --- | --- | --- |
| *Predictors* | *Log-Odds* | *std. Error* | *CI* | *p* |
| (Intercept) | -3.56 | 1.01 | -6.43 – -2.02 | - |
| Habitat [Urban] | 1.48 | 1.10 | -0.35 – 4.44 | 0.123 |
| Observations | 90 | | | |

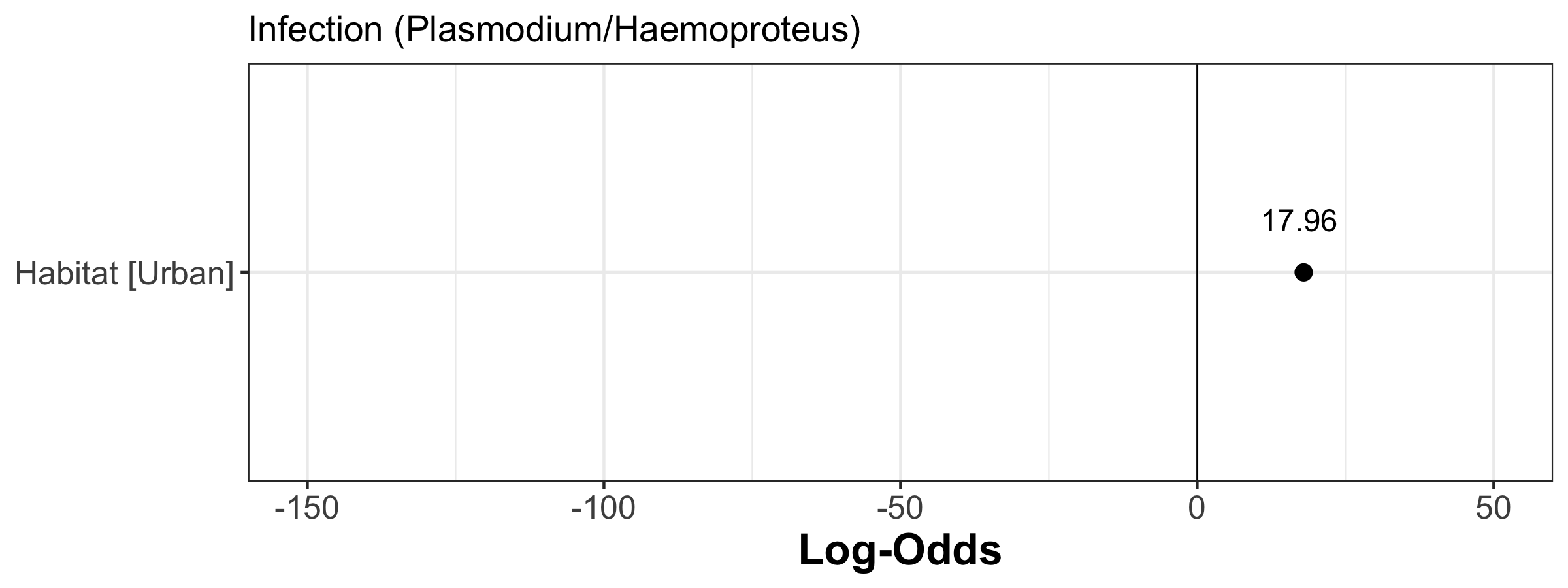

**Figure S3:** Forest plot of the estimates (Log-odds) of the model investigating the factors associated with Plasmodium/Haemoproteus infection using habitat as a binary variable (urban vs. forest) in nestlings | The estimate value is indicated by plain points and its associate 95% confidence intervals by plain segments.

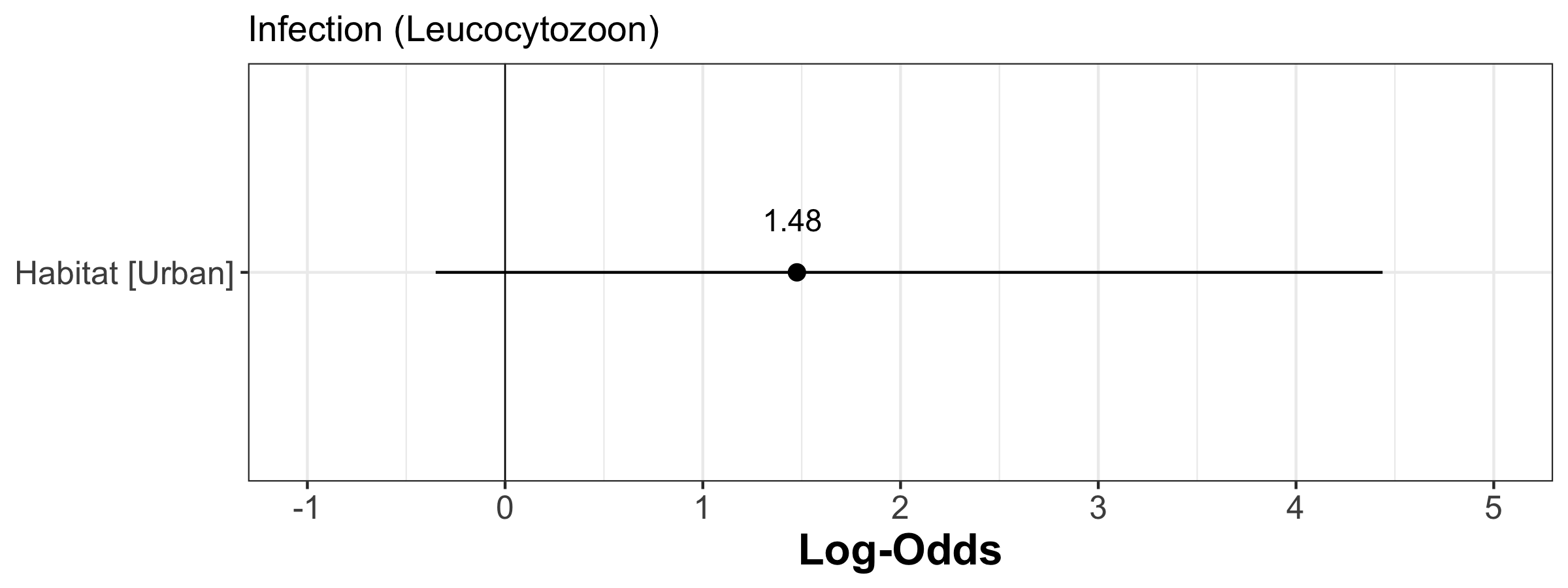

**Figure S4:** Forest plot of the estimates (Log-odds) of the model investigating the factors associated with Leucocytozoon infection using habitat as a binary variable (urban vs. forest) in nestlings | The estimate value is indicated by plain points and its associate 95% confidence intervals by plain segments.

##### 4.1.2. Linear model assumptions

###### 4.1.2.1. Distribution of residuals

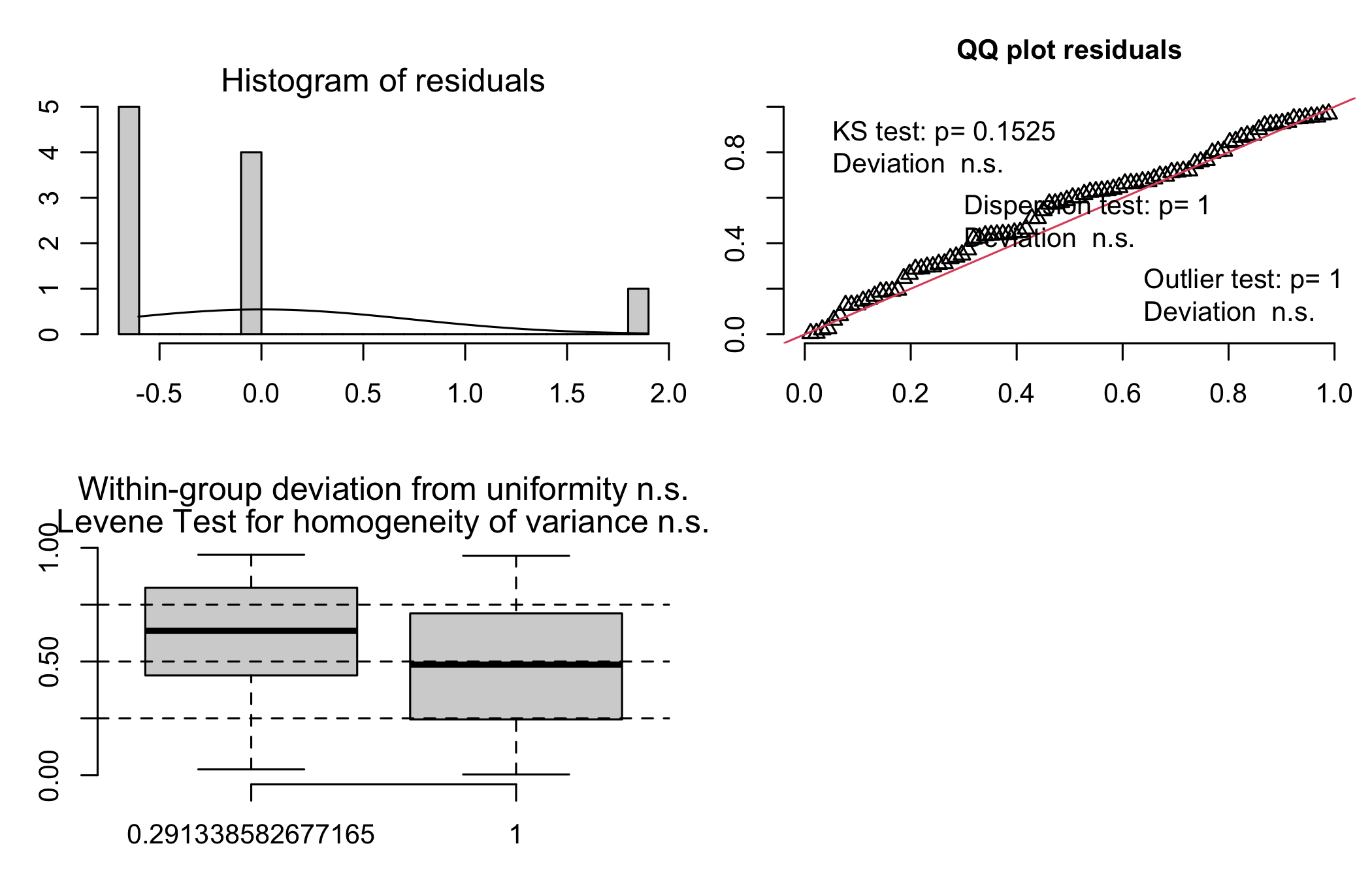

**Figure S5:** Model assumption check of the model on Plasmodium/Haemoproteus infection probability with habitat as a binary variable (urban vs. forest) in nestlings | Depicted are the histogram of residuals, the Q-Q plot, and the scatter plot of the fitted values vs the residuals.

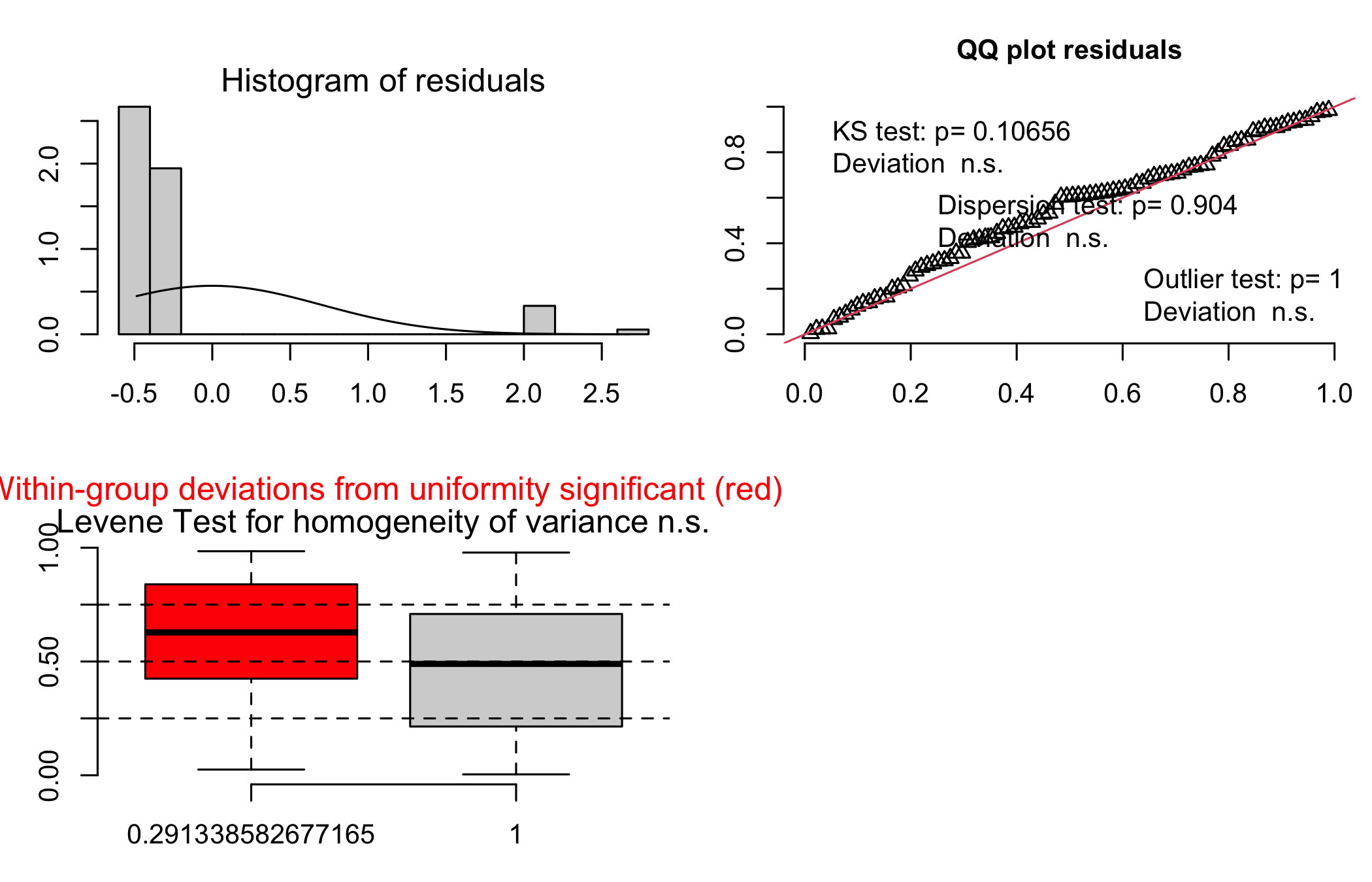

**Figure S6:** Model assumption check of the model on Leucocytozoon infection probability with habitat as a binary variable (urban vs. forest) in nestlings | Depicted are the histogram of residuals, the Q-Q plot, and the scatter plot of the fitted values vs the residuals. Despite statistical significance (as the tests are highly sensitive, see DHARMa package vignette), deviations appeared reasonable and inconsequential (Schielzeth et al. 2020), as strengthened by further investigation (see below).

##### 4.1.3. Linear model stability

**Table S5:** Dfbetas of the model with habitat as a binary variable (urban vs. forest) on Plasmodium/Haemoproteus infection probability in nestlings.

| **Estimate** | **Min** | **Max** |
| --- | --- | --- |
| -19.57 | -19.59 | -19.57 |
| 17.96 | 17.93 | 18.05 |

**Table S6:** Dfbetas of the model with habitat as a binary variable (urban vs. forest) on Leucocytozoon infection probability in nestlings.

| **Estimate** | **Min** | **Max** |
| --- | --- | --- |
| -3.56 | -3.60 | -3.09 |
| 1.48 | 1.01 | 1.60 |

#### 4.2. Urbanization: averaged urbanization level by site

##### 4.2.1. Linear model results

**Table S7:** Detailed output of fixed effects included in the GLM investigating *Plasmodium/Haemoproteus* prevalence in nestlings with urbanization level averaged per site.

|  | **Infection**  **(Plasmodium/Haemoproteus)** | | | |
| --- | --- | --- | --- | --- |
| *Predictors* | *Log-Odds* | *std. Error* | *CI* | *p* |
| (Intercept) | -2.20 | 0.36 | -2.98 – -1.56 | **-** |
| Site-level naturalness | -0.29 | 0.26 | -0.79 – 0.25 | 0.276 |
| Observations | 90 | | | |

**Table S8:** Detailed output of fixed effects included in the GLM investigating *Leucocytozoon* prevalence in nestlings with urbanization level averaged per site.

|  | **Infection (Leucocytozoon)** | | | |
| --- | --- | --- | --- | --- |
| *Predictors* | *Log-Odds* | *std. Error* | *CI* | *p* |
| (Intercept) | -2.49 | 0.41 | -3.40 – -1.78 | **-** |
| Site-level naturalness | -0.33 | 0.28 | -0.89 – 0.26 | 0.256 |
| Observations | 90 | | | |

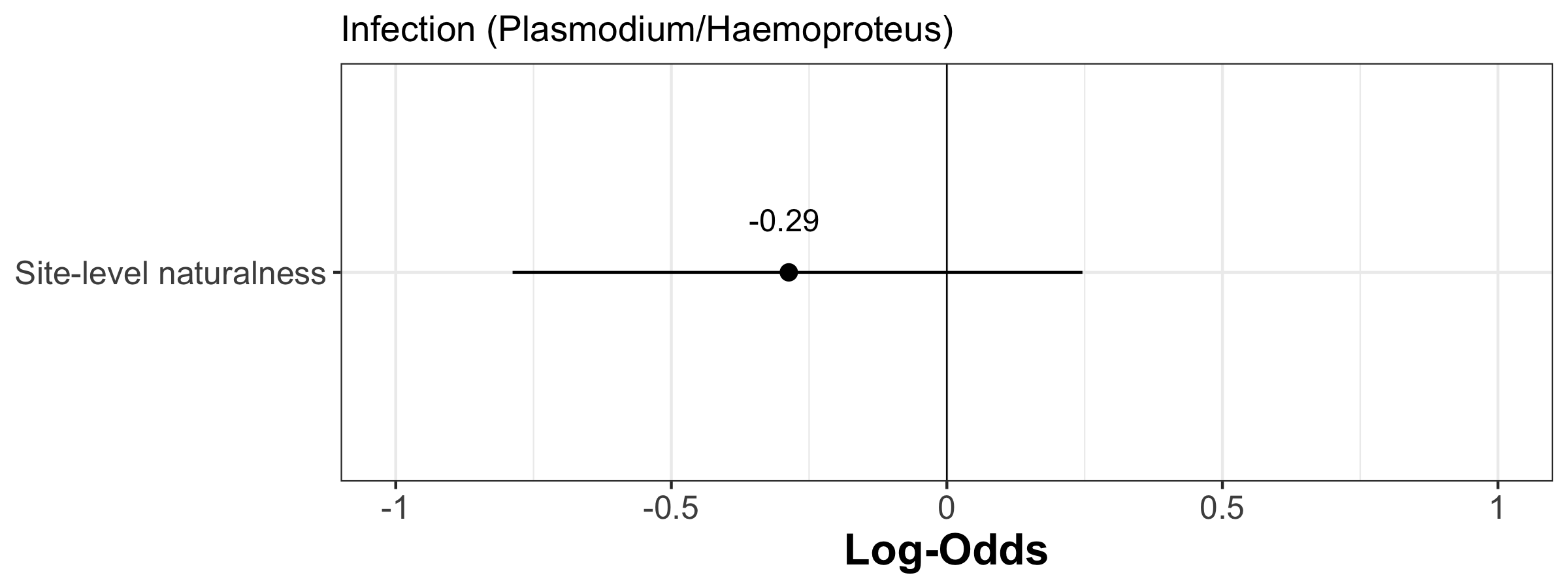

**Figure S7:** Forest plot of the estimates (Log-odds) of the model investigating the factors associated with Plasmodium/Haemoproteus infection using urbanization level averaged per site in nestlings | The estimate value is indicated by plain points and its associate 95% confidence intervals by plain segments.

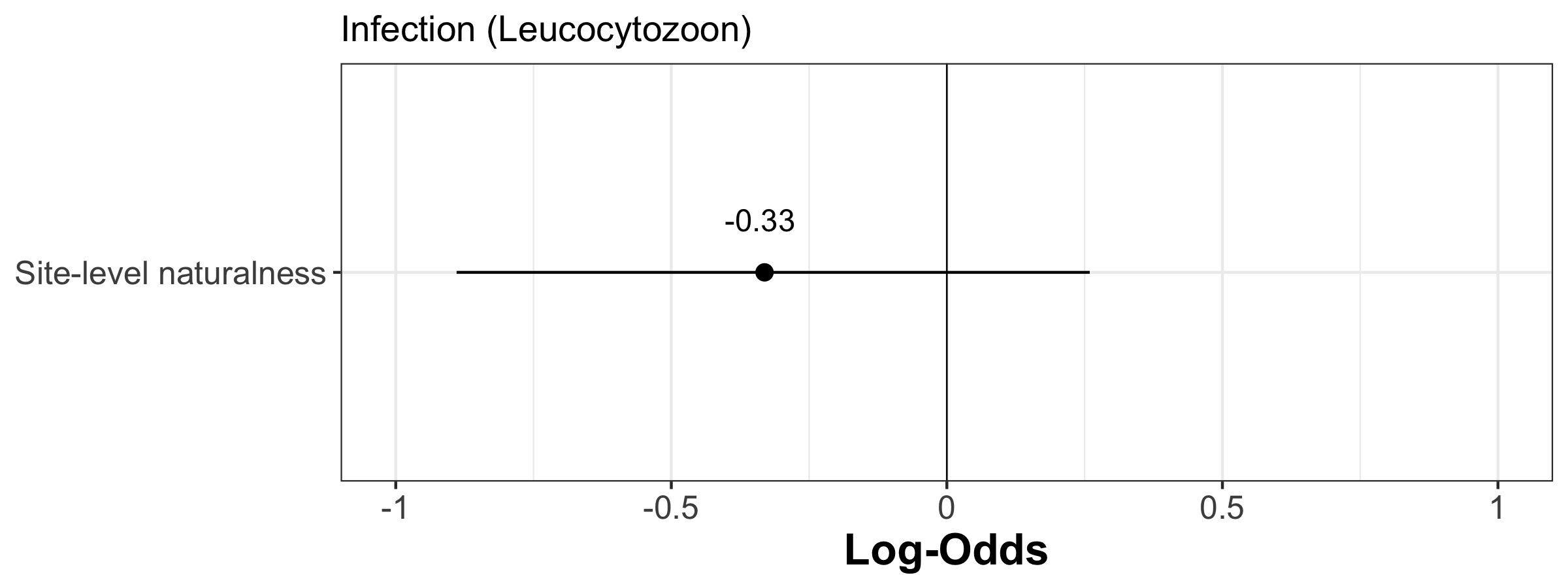

**Figure S8:** Forest plot of the estimates (Log-odds) of the model investigating the factors associated with Leucocytozoon infection using urbanization level averaged per site in nestlings | The estimate value is indicated by plain points and its associate 95% confidence intervals by plain segments.

##### 4.2.2. Linear model assumptions

###### 4.2.2.1. Distribution of residuals

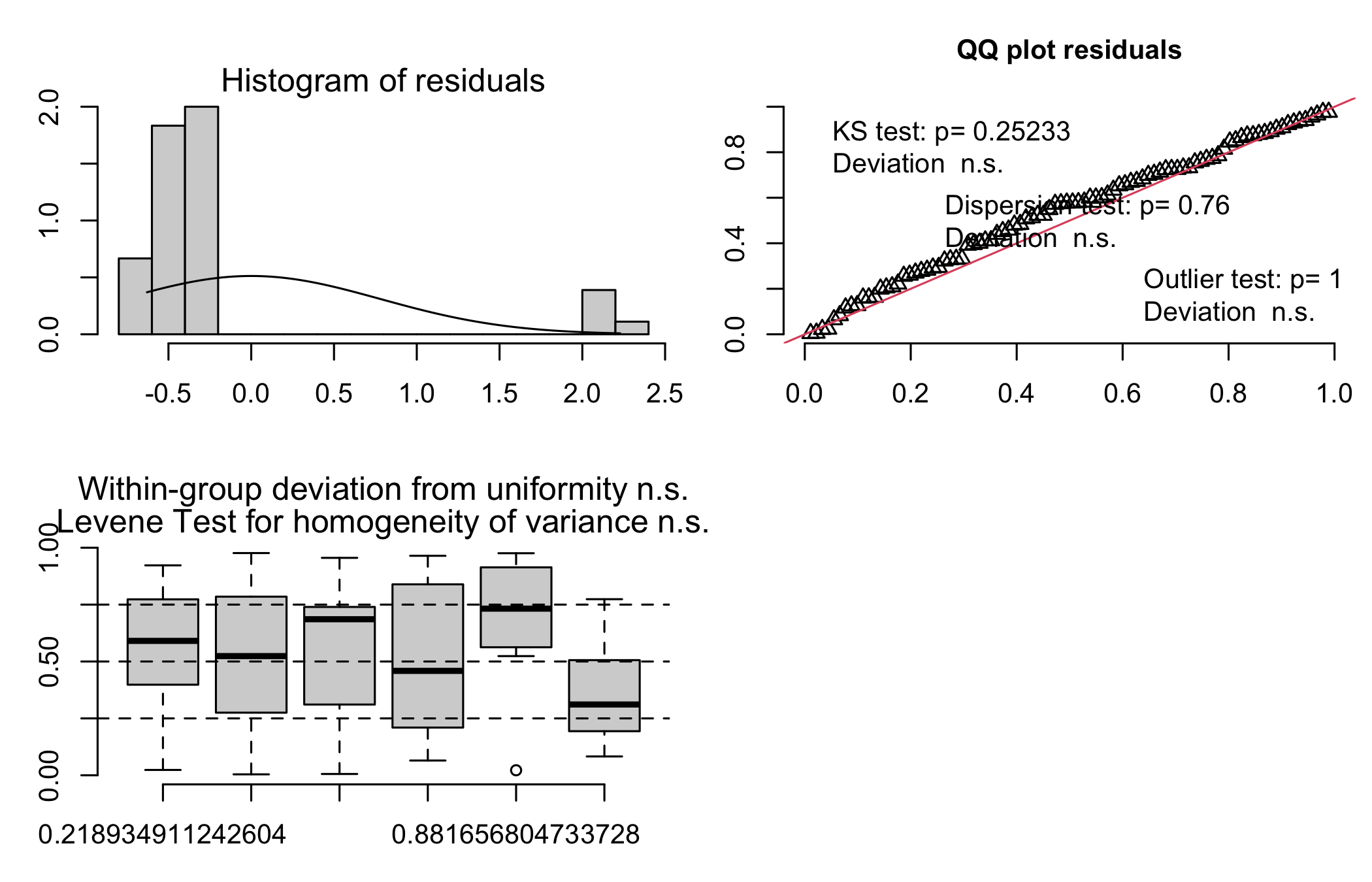

**Figure S9**: Model assumption check of the model on Plasmodium/Haemoproteus infection probability with urbanization level averaged per site in nestlings | Depicted are the histogram of residuals, the Q-Q plot, and the scatter plot of the fitted values vs the residuals.

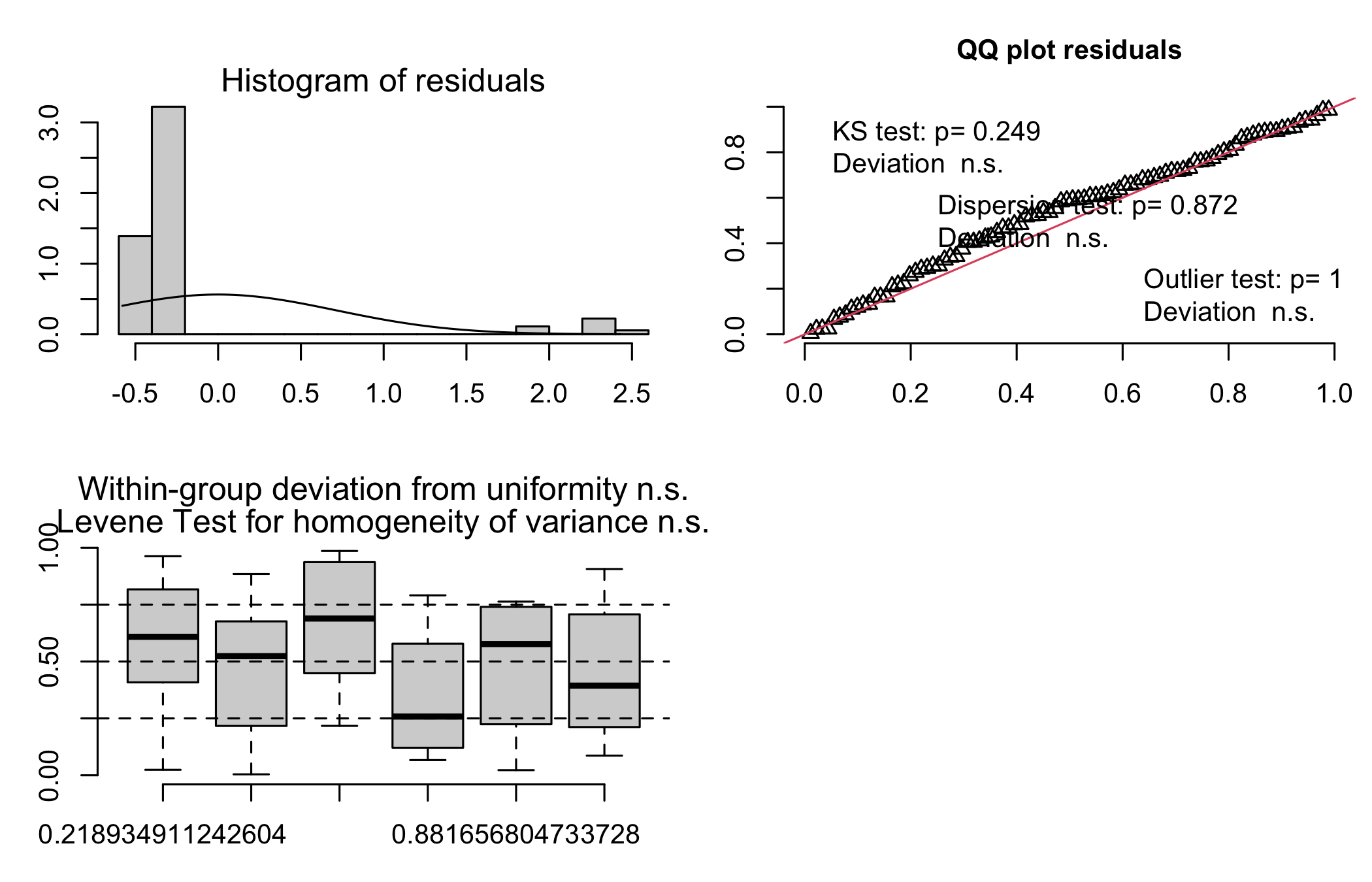

**Figure S10:** Model assumption check of the model on Leucocytozoon infection probability with urbanization level averaged per site in nestlings | Depicted are the histogram of residuals, the Q-Q plot, and the scatter plot of the fitted values vs the residuals.

##### 4.2.3. Linear model stability

**Table S9:** Dfbetas of the model with urbanization level averaged per site on Plasmodium/Haemoproteus infection probability in nestlings.

| **Estimate** | **Min** | **Max** |
| --- | --- | --- |
| -2.20 | -2.22 | -2.11 |
| -0.29 | -0.32 | -0.25 |

**Table S10:** Dfbetas of the model with urbanization level averaged per site on Leucocytozoon infection probability in nestlings.

| **Estimate** | **Min** | **Max** |
| --- | --- | --- |
| -2.49 | -2.52 | -2.38 |
| -0.33 | -0.46 | -0.26 |

#### 4.3. Urbanization as a gradient

##### 4.3.1. Linear model results

**Table S11:** Detailed output of fixed effects included in the GLM investigating *Plasmodium/Haemoproteus* prevalence in nestlings with urbanization level per nest-box.

|  | **Infection (Plasmodium/Haemoproteus)** | | | |
| --- | --- | --- | --- | --- |
| *Predictors* | *Log-Odds* | *std. Error* | *CI* | *p* |
| (Intercept) | -2.19 | 0.36 | -2.97 – -1.55 | - |
| Nest-level naturalness | -0.03 | 0.28 | -0.55 – 0.58 | 0.908 |
| Observations | 90 | | | |

**Table S12:** Detailed output of fixed effects included in the GLM investigating *Leucocytozoon* prevalence in nestlings with urbanization level averaged per nest-box.

|  | **Infection (Leucocytozoon)** | | | |
| --- | --- | --- | --- | --- |
| *Predictors* | *Log-Odds* | *std. Error* | *CI* | *p* |
| (Intercept) | -2.48 | 0.41 | -3.39 – -1.76 | - |
| Nest-level naturalness | -0.39 | 0.28 | -0.96 – 0.19 | 0.175 |
| Observations | 90 | | | |

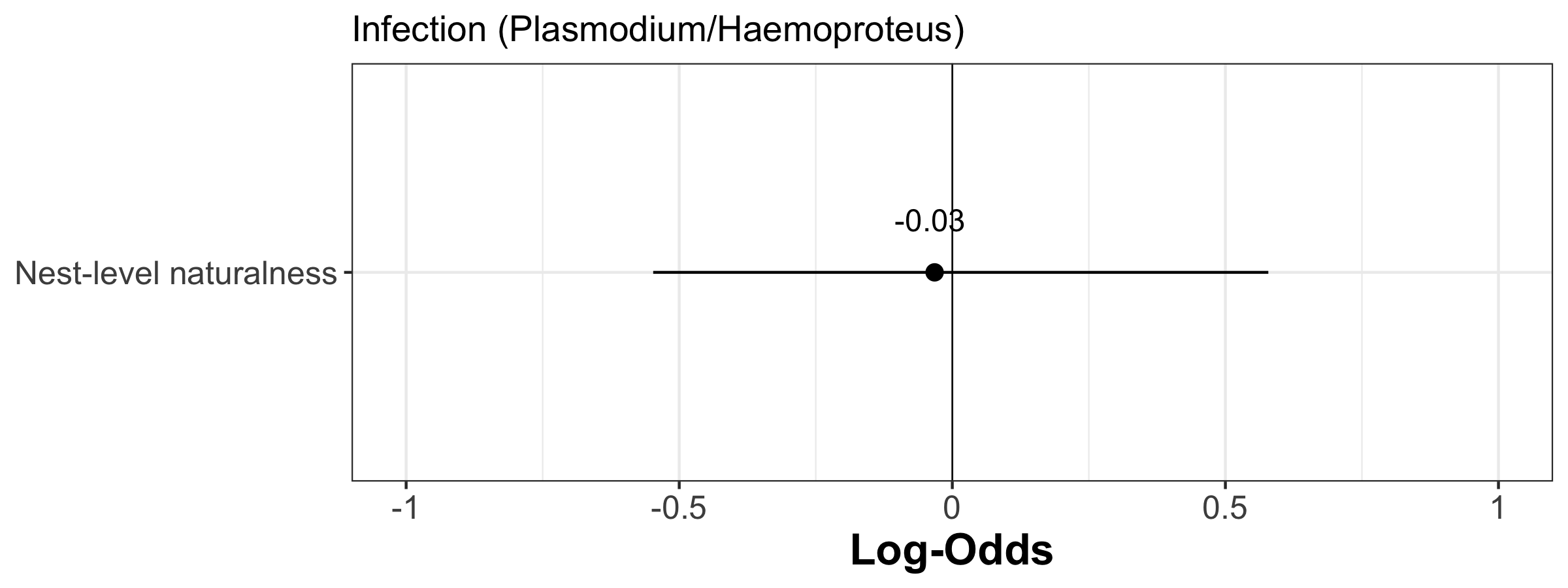

**Figure S11:** Forest plot of the estimates (Log-odds) of the model investigating the factors associated with Plasmodium/Haemoproteus infection using urbanization gradient at the nest-box level in nestlings | The estimate value is indicated by plain points and its associate 95% confidence intervals by plain segments.

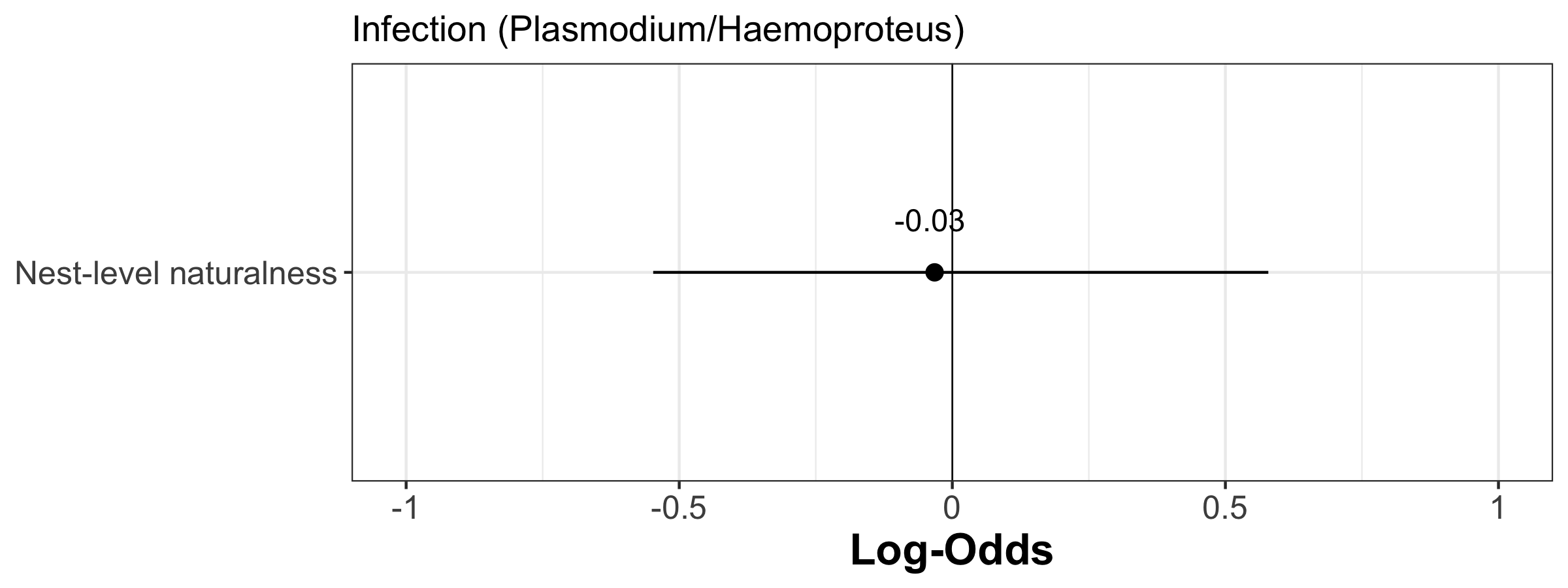

**Figure S12:** Forest plot of the estimates (Log-odds) of the model investigating the factors associated with Leucocytozoon infection using urbanization gradient at the nest-box level in nestlings | The estimate value is indicated by plain points and its associate 95% confidence intervals by plain segments.

##### 4.3.2. Linear model assumptions

###### Distribution of residuals

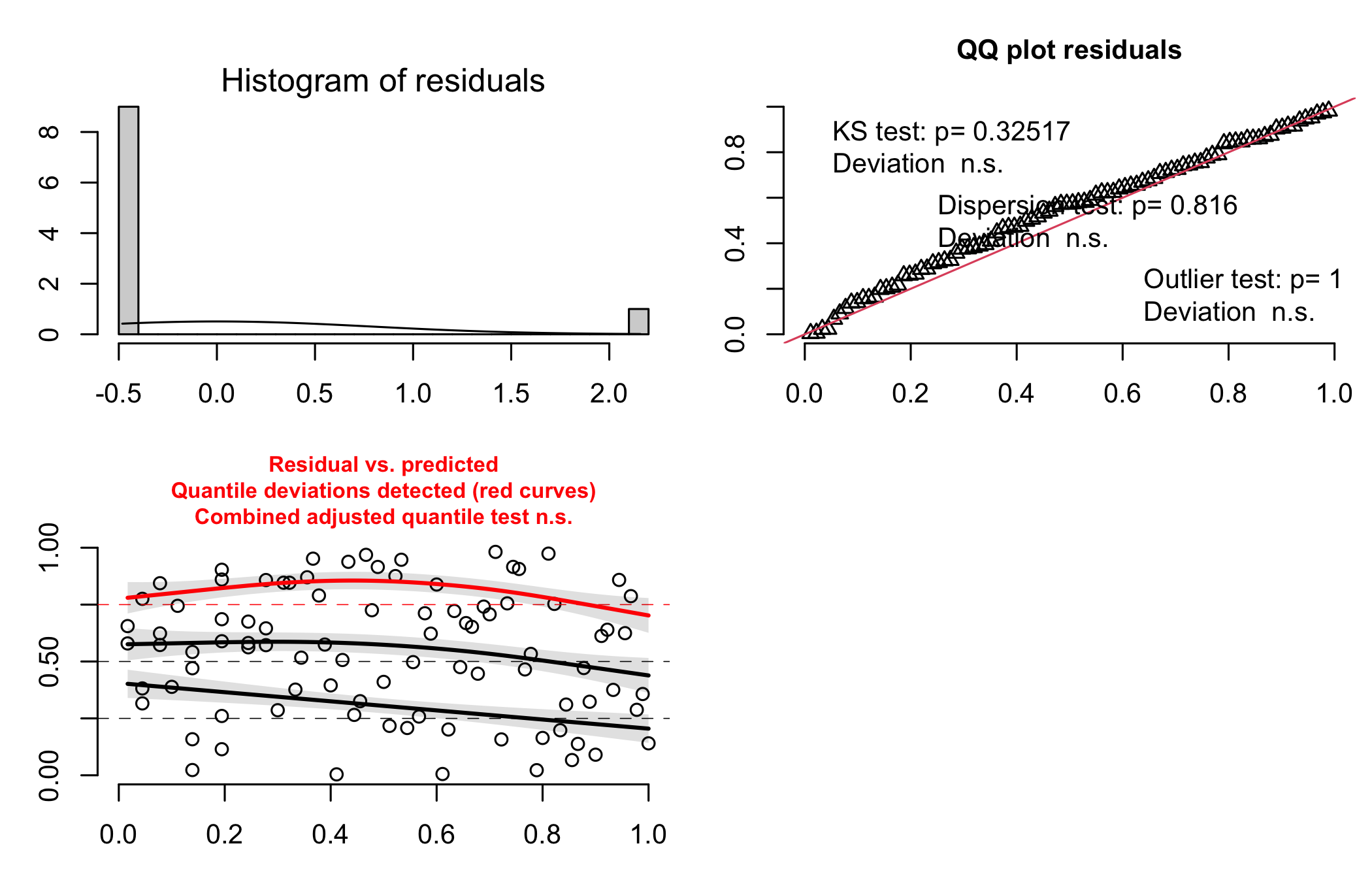

**Figure S13:** Model assumption check of the model on Plasmodium/Haemoproteus infection probability with the urbanization gradient at the nest-box level in nestlings | Depicted are the histogram of residuals, the Q-Q plot, and the scatter plot of the fitted values vs the residuals.

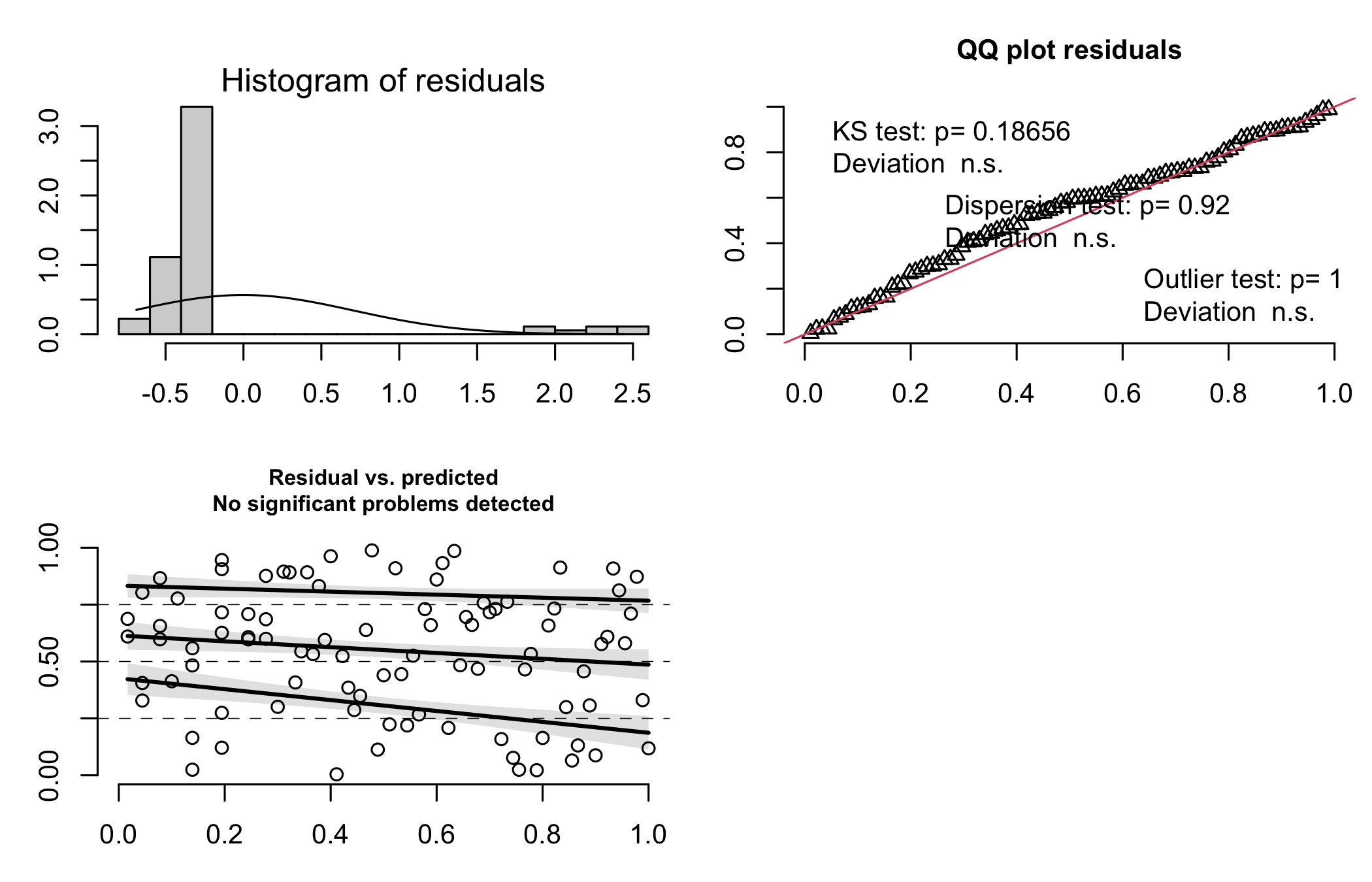

**Figure S14:** Model assumption check of the model on Leucocytozoon infection probability with urbanization gradient at the nest-box level in nestlings | Depicted are the histogram of residuals, the Q-Q plot, and the scatter plot of the fitted values vs the residuals.

##### 4.3.3. Linear model stability

**Table S13:** Dfbetas of the model with the urbanization gradient at the nest-box level on Plasmodium/Haemoproteus infection probability in nestlings

| **Estimate** | **Min** | **Max** |
| --- | --- | --- |
| -2.19 | -2.22 | -2.09 |
| -0.03 | -0.11 | 0.02 |

**Table S14:** Dfbetas of the model with the urbanization gradient at the nest-box level on Leucocytozoon infection probability in nestlings

| **Estimate** | **Min** | **Max** |
| --- | --- | --- |
| -2.48 | -2.51 | -2.37 |
| -0.39 | -0.52 | -0.32 |

### 5. Adult models

#### 5.1. Urbanization as binary variable (urban vs. rural)

###### 5.1.1. Linear model results

**Table S15:** Detailed output of fixed effects included in the GLM investigating *Plasmodium/Haemoproteus* prevalence in adults with urbanization as a binary variable (urban vs. forest).

|  | **Infection (Plasmodium/Haemoproteus)** | | | |
| --- | --- | --- | --- | --- |
| *Predictors* | *Log-Odds* | *std. Error* | *CI* | *p* |
| (Intercept) | 4.10 | 1.21 | 2.03 – 7.18 | - |
| Habitat [Urban] | -0.07 | 1.18 | -3.10 – 2.01 | 0.954 |
| Age | -0.31 | 0.34 | -0.94 – 0.42 | 0.379 |
| Sex [Male] | 0.73 | 0.73 | -0.65 – 2.31 | 0.305 |
| Year [2019] | -0.73 | 0.79 | -2.48 – 0.75 | 0.344 |
| Observations | 267 | | | |

**Table S16:** Detailed output of fixed effects included in the GLM investigating *Leucocytozoon* prevalence in adults with urbanization as a binary variable (urban vs. forest).

|  | **Infection (Leucocytozoon)** | | | |
| --- | --- | --- | --- | --- |
| *Predictors* | *Log-Odds* | *std. Error* | *CI* | *p* |
| (Intercept) | 1.92 | 0.72 | 0.54 – 3.43 | - |
| Habitat [Urban] | -0.75 | 0.60 | -2.06 – 0.35 | 0.191 |
| Age | 0.27 | 0.30 | -0.28 – 0.89 | 0.351 |
| Sex [Male] | 0.23 | 0.43 | -0.62 – 1.09 | 0.591 |
| Year [2019] | 1.05 | 0.48 | 0.13 – 2.06 | **0.024** |
| Observations | 267 | | | |

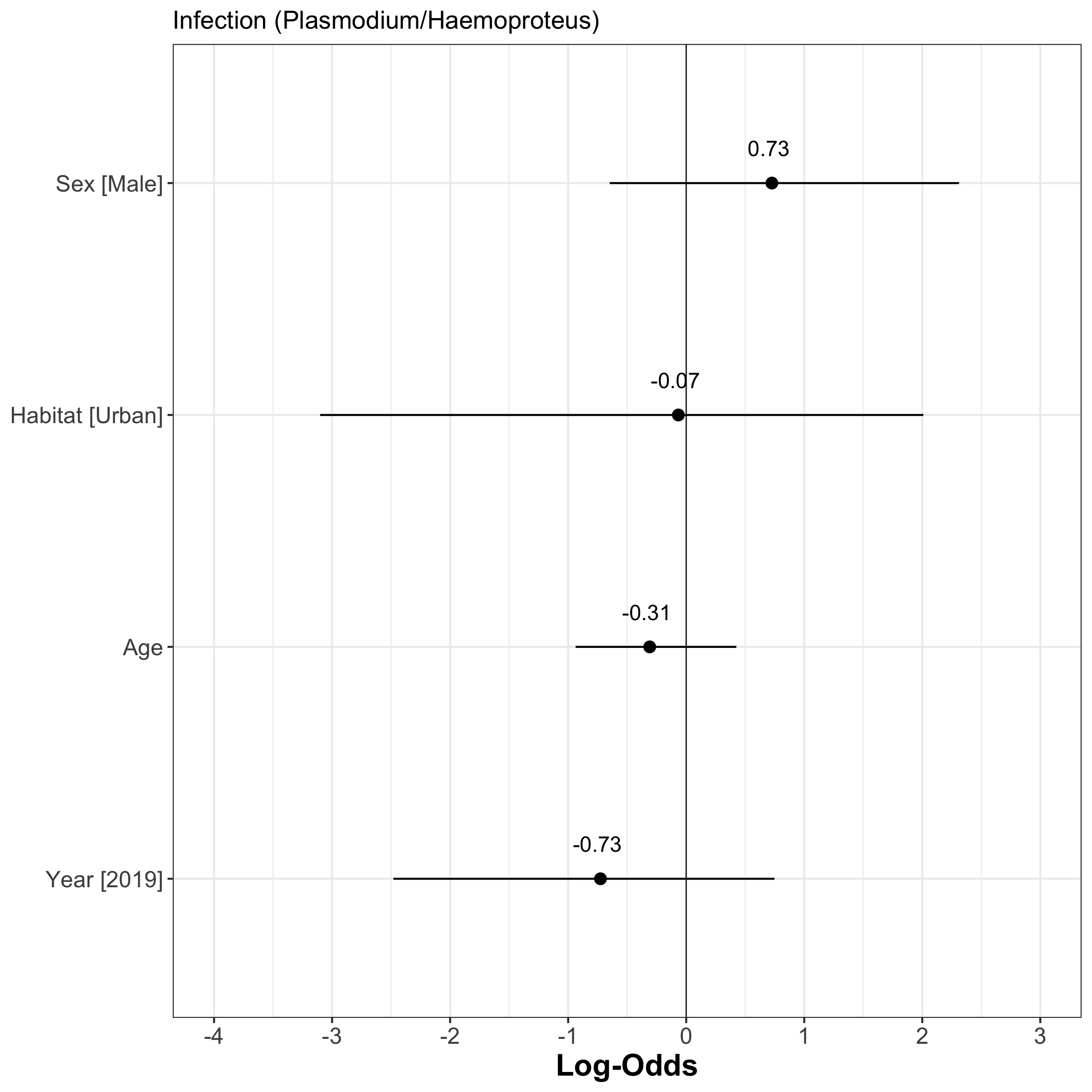

**Figure S15:** Forest plot of the estimates (Log-odds) of the model investigating the factors associated with Plasmodium/Haemoproteus infection using habitat as a binary variable (urban vs. forest) | The estimate value is indicated by plain points and its associate 95% confidence intervals by plain segments.

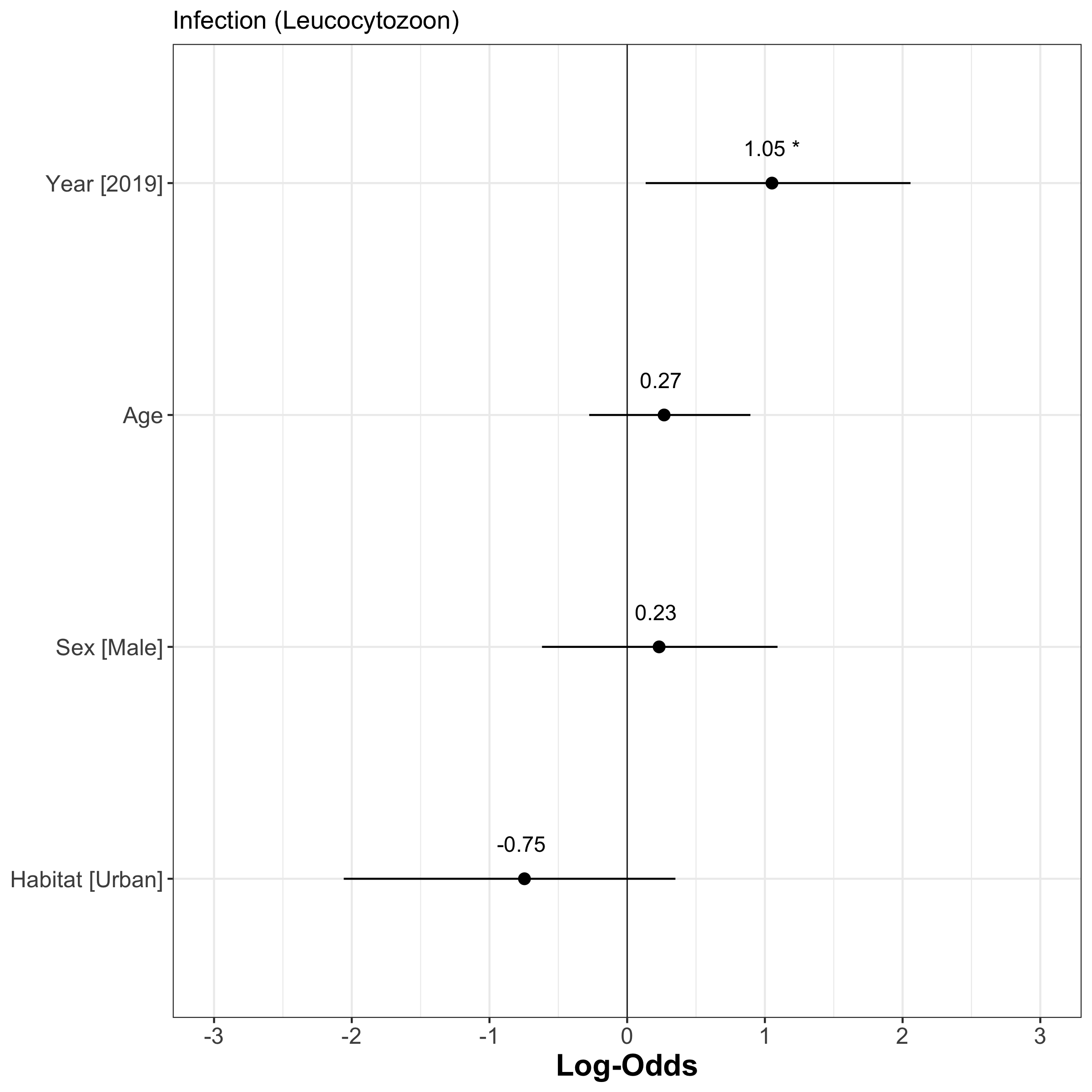

**Figure S16:** Forest plot of the estimates (Log-odds) of the model investigating the factors associated with Leucocytozoon infection using habitat as a binary variable (urban vs. forest) | The estimate value is indicated by plain points and its associate 95% confidence intervals by plain segments.

##### 5.1.2. Linear model assumptions

###### 5.1.2.1. Collinearity

**Table S17:** Collinearity assessment of the model investigating the factors associated with Plasmodium/Haemoproteus infection using habitat as a binary variable (urban vs. forest) | VIF = Variance Inflation Factor, CI = Confidence interval, SE = Standard Error. This assessment is based on the 'check_collinarity' function of the performance package.

| **Term** | **VIF** | **VIF_CI_low** | **VIF_CI_high** | **SE_factor** | **Tolerance** | **Tolerance**  **CI_low** | **Tolerance**  **CI_high** |
| --- | --- | --- | --- | --- | --- | --- | --- |
| **Habitat** | 1.41 | 1.24 | 1.68 | 1.19 | 0.71 | 0.59 | 0.80 |
| **Site-level naturalness** | 2.23 | 1.89 | 2.70 | 1.49 | 0.45 | 0.37 | 0.53 |
| **Nest-level naturalness** | 1.81 | 1.55 | 2.17 | 1.34 | 0.55 | 0.46 | 0.64 |
| **Sex** | 1.03 | 1.00 | 2.49 | 1.02 | 0.97 | 0.40 | 1.00 |
| **Age** | 1.08 | 1.01 | 1.43 | 1.04 | 0.93 | 0.70 | 0.99 |
| **Year** | 1.36 | 1.21 | 1.63 | 1.17 | 0.73 | 0.61 | 0.83 |

**Table S18:** Collinearity assessment of the model investigating the factors associated with Leucocytozoon infection using habitat as a binary variable (urban vs. forest) | VIF = Variance Inflation Factor, CI = Confidence interval, SE = Standard Error. This assessment is based on the 'check_collinarity' function of the performance package.

| **Term** | **VIF** | **VIF_CI_low** | **VIF_CI_high** | **SE_factor** | **Tolerance** | **Tolerance**  **CI_low** | **Tolerance_**  **CI_high** |
| --- | --- | --- | --- | --- | --- | --- | --- |
| **Habitat** | 1.42 | 1.25 | 1.70 | 1.19 | 0.71 | 0.59 | 0.80 |
| **Site-level naturalness** | 1.47 | 1.29 | 1.75 | 1.21 | 0.68 | 0.57 | 0.78 |
| **Age** | 1.05 | 1.00 | 1.67 | 1.02 | 0.95 | 0.60 | 1.00 |
| **Sex** | 1.03 | 1.00 | 2.89 | 1.01 | 0.97 | 0.35 | 1.00 |
| **Year** | 1.15 | 1.05 | 1.41 | 1.07 | 0.87 | 0.71 | 0.95 |

###### 5.1.2.2. Distribution of residuals

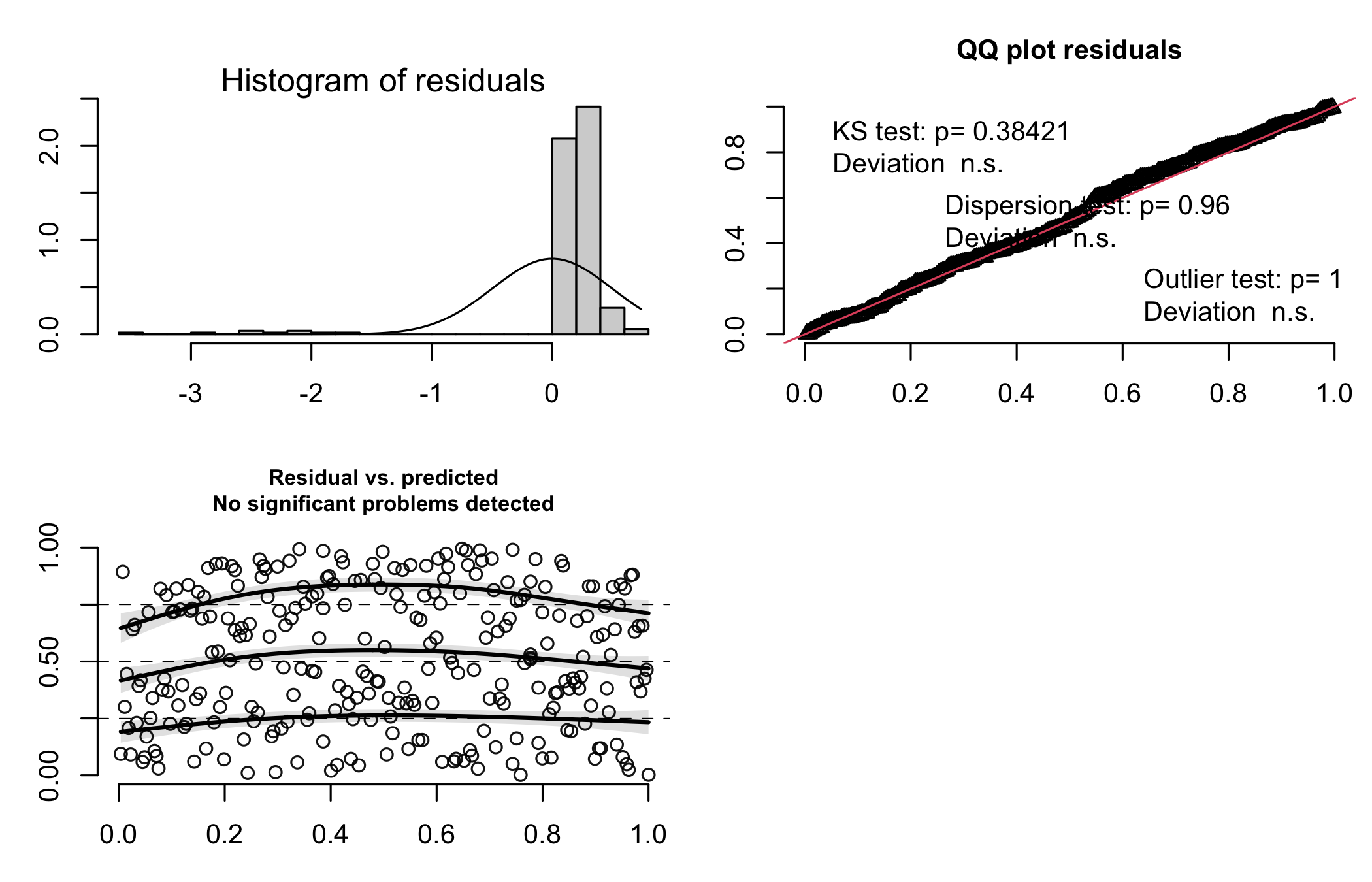

**Figure S17:** Model assumption check of the model on Plasmodium/Haemoproteus infection probability with habitat as a binary variable (urban vs. forest) | Depicted are the histogram of residuals, the Q-Q plot, and the scatter plot of the fitted values vs the residuals.

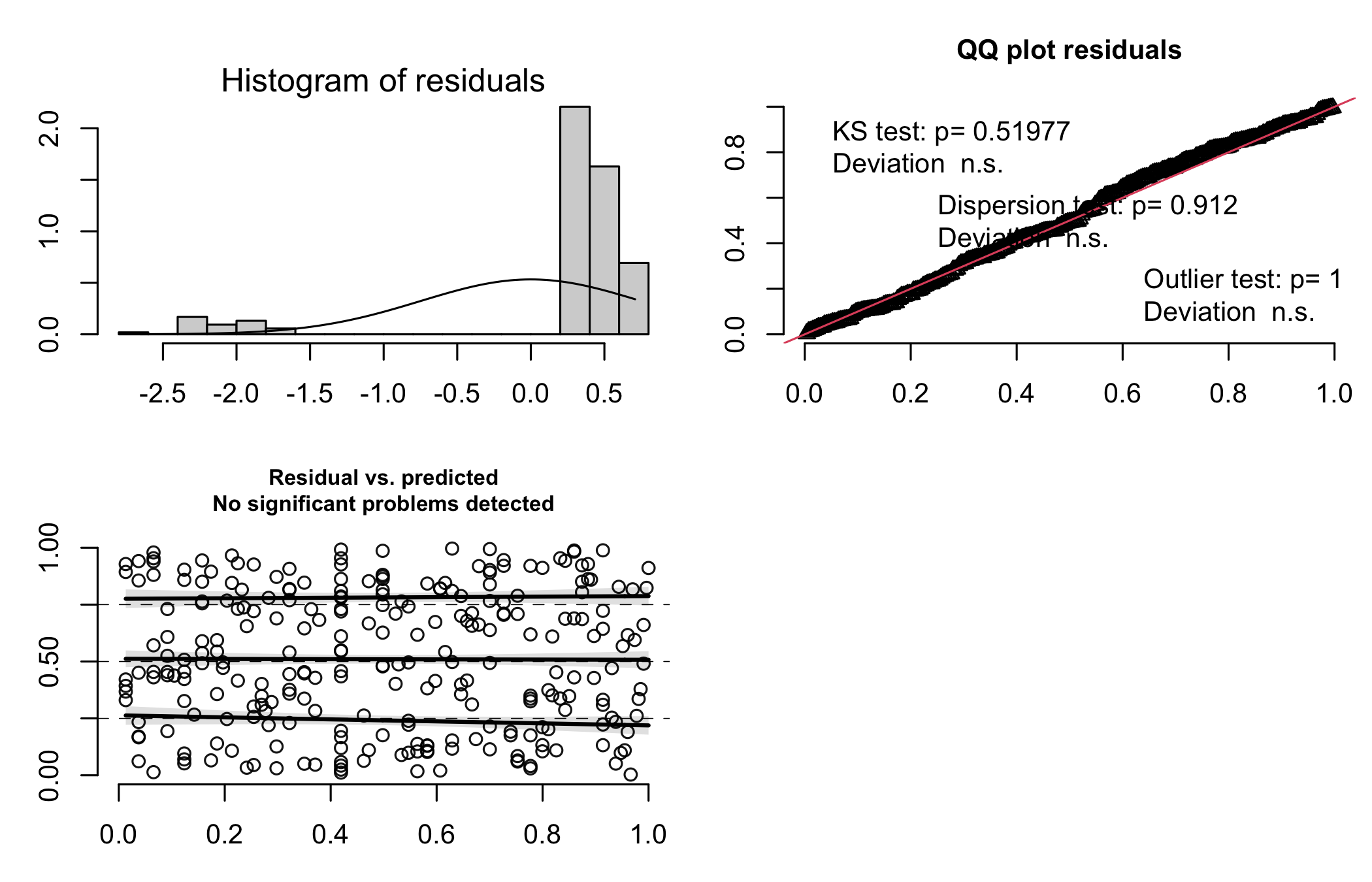

**Figure S18:** Model assumption check of the model on Leucocytozoon infection probability with habitat as a binary variable (urban vs. forest) | Depicted are the histogram of residuals, the Q-Q plot, and the scatter plot of the fitted values vs the residuals.

##### 5.1.3. Linear model stability

**Table S19:** Dfbetas of the model with habitat as a binary variable (urban vs. forest) on Plasmodium/Haemoproteus infection probability.

| **Estimate** | **Min** | **Max** |
| --- | --- | --- |
| 4.78 | 4.46 | 5.83 |
| -0.52 | -0.77 | -0.21 |
| 0.32 | 0.12 | 0.45 |
| -0.87 | -1.04 | -0.76 |
| 0.66 | 0.46 | 0.82 |
| -0.30 | -0.77 | -0.18 |
| -0.97 | -1.21 | -0.76 |

**Table S20:** Dfbetas of the model with habitat as a binary variable (urban vs. forest) on Leucocytozoon infection probability.

| **Estimate** | **Min** | **Max** |
| --- | --- | --- |
| 2.21 | 1.98 | 2.50 |
| -1.06 | -1.18 | -0.90 |
| -0.23 | -0.27 | -0.13 |
| 0.26 | 0.08 | 0.31 |
| 0.22 | 0.13 | 0.30 |
| 0.95 | 0.83 | 1.05 |

#### 5.2. Urbanization: averaged urbanization level by site

##### 5.2.1. Linear model results

**Table S21:** Detailed output of fixed effects included in the GLM investigating *Plasmodium/Haemoproteus* prevalence in adults with urbanization level averaged per site.

|  | **Infection (Plasmodium/Haemoproteus)** | | | |
| --- | --- | --- | --- | --- |
| *Predictors* | *Log-Odds* | *std. Error* | *CI* | *p* |
| (Intercept) | 4.11 | 0.90 | 2.41 – 6.03 | **-** |
| Site-level naturalness | -0.21 | 0.36 | -1.01 – 0.44 | 0.548 |
| Age | -0.31 | 0.33 | -0.93 – 0.42 | 0.377 |
| Sex [Male] | 0.73 | 0.73 | -0.65 – 2.32 | 0.302 |
| Year [2019] | -0.94 | 0.81 | -2.68 – 0.59 | 0.234 |
| Observations | 267 | | | |

**Table S22:** Detailed output of fixed effects included in the GLM investigating *Leucocytozoon* prevalence in adults with urbanization level averaged per site.

|  | Infection (Leucocytozoon) | | | |
| --- | --- | --- | --- | --- |
| *Predictors* | *Log-Odds* | *std. Error* | *CI* | *p* |
| (Intercept) | 1.47 | 0.60 | 0.28 – 2.64 | - |
| Site-level naturalness | -0.02 | 0.20 | -0.44 – 0.35 | 0.911 |
| Age | 0.21 | 0.29 | -0.33 – 0.84 | 0.500 |
| Sex [Male] | 0.26 | 0.43 | -0.59 – 1.12 | 0.548 |
| Year [2019] | 0.84 | 0.50 | -0.12 – 1.87 | 0.087 |
| Observations | 267 | | | |

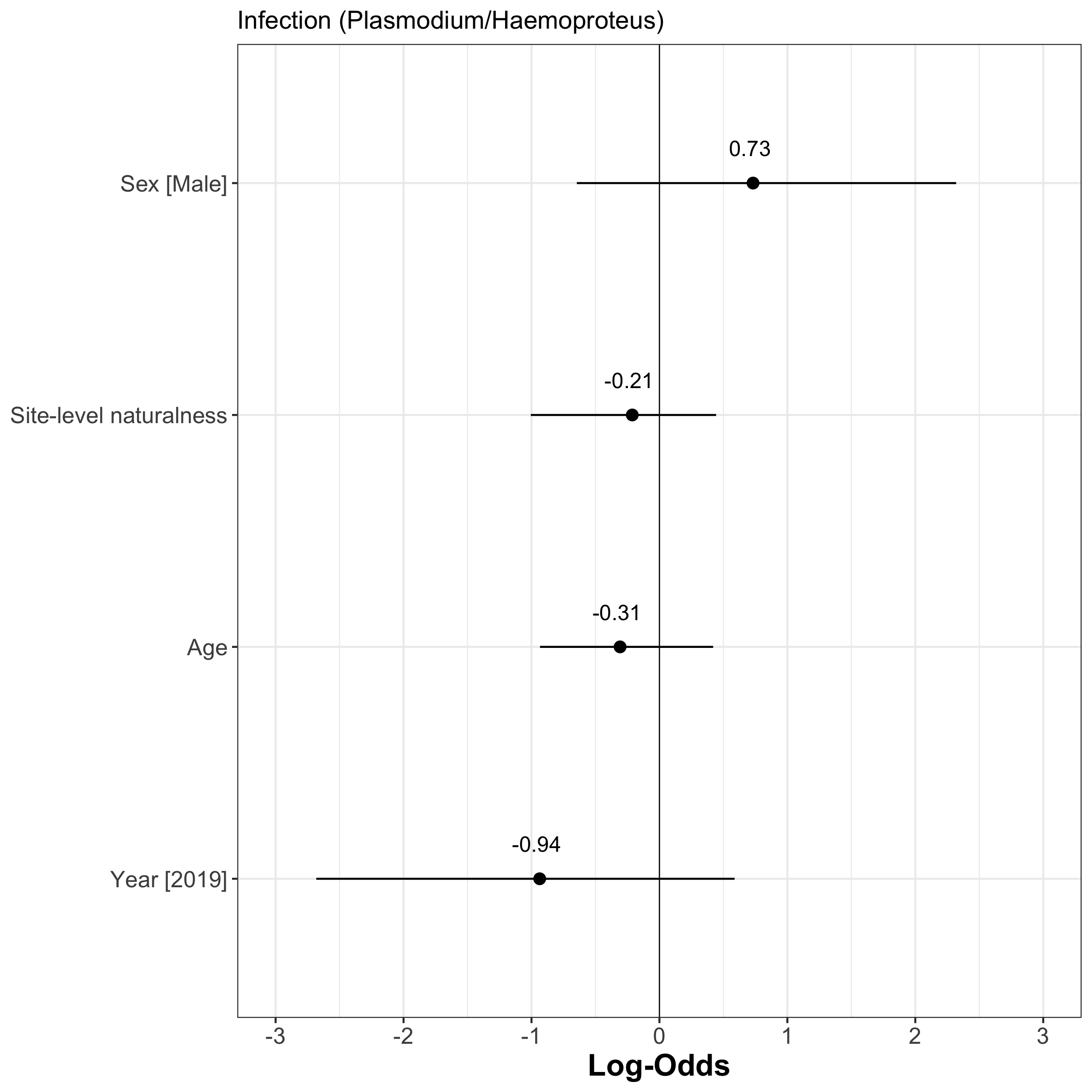

**Figure S19:** Forest plot of the estimates (Log-odds) of the model investigating the factors associated with Plasmodium/Haemoproteus infection using urbanization level averaged per site | The estimate value is indicated by plain points and its associate 95% confidence intervals by plain segments.

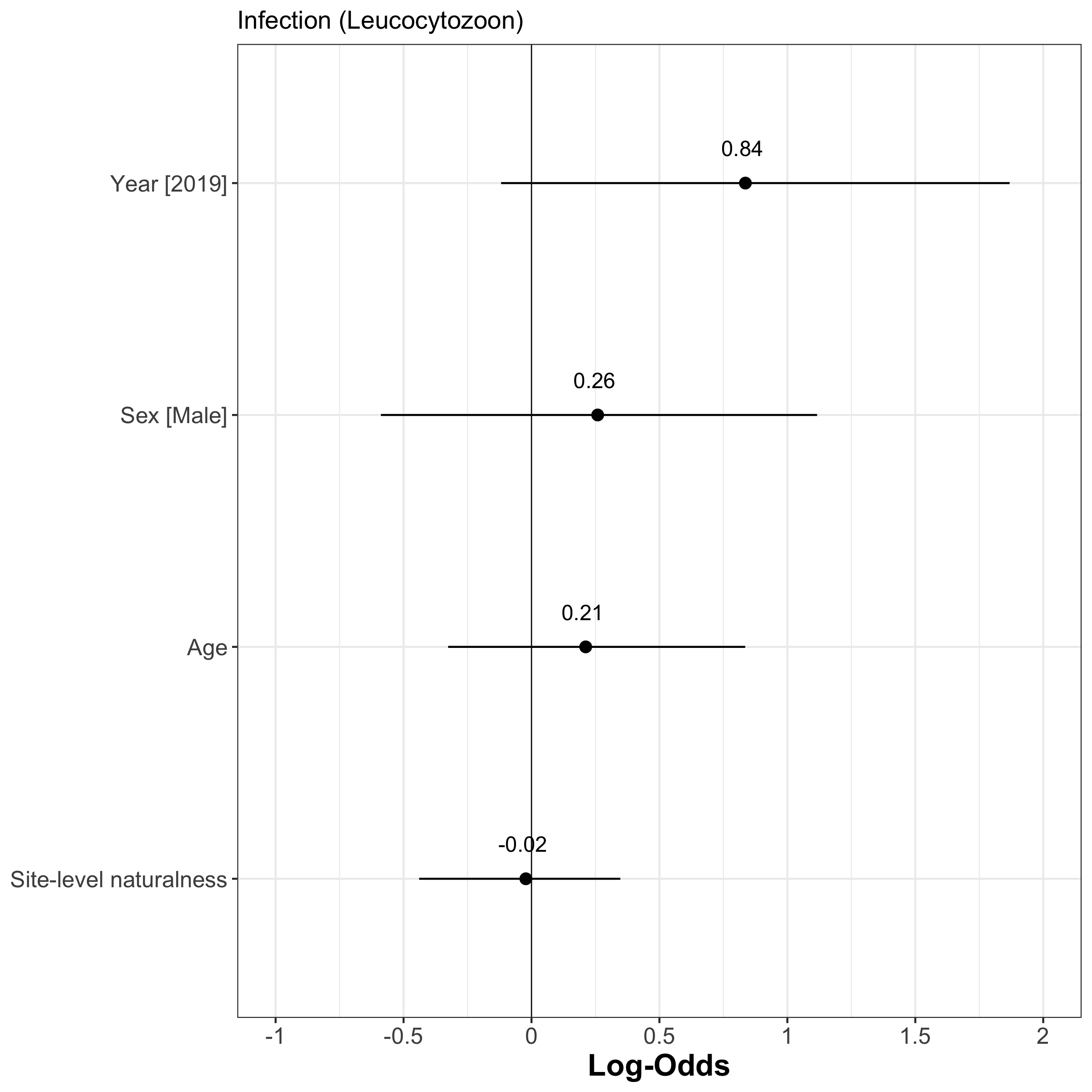

**Figure S20:** Forest plot of the estimates (Log-odds) of the model investigating the factors associated with Leucocytozoon infection using urbanization level averaged per site| The estimate value is indicated by plain points and its associate 95% confidence intervals by plain segments.

##### 5.2.2. Linear model assumptions

###### 5.2.2.1. Collinearity

**Table S23:** Collinearity assessment of the model investigating the factors associated with Plasmodium/Haemoproteus infection using urbanization level averaged per site | VIF = Variance Inflation Factor, CI = Confidence interval, SE = Standard Error. This assessment is based on the 'check_collinarity' function of the performance package.

| **Term** | **VIF** | **VIF_CI_low** | **VIF_CI_high** | **SE_factor** | **Tolerance** | **Tolerance**  **CI_low** | **Tolerance**  **CI_high** |
| --- | --- | --- | --- | --- | --- | --- | --- |
| **Site-level naturalness** | 1.23 | 1.11 | 1.48 | 1.11 | 0.82 | 0.68 | 0.90 |
| **Age** | 1.06 | 1.01 | 1.53 | 1.03 | 0.94 | 0.65 | 0.99 |
| **Sex** | 1.02 | 1.00 | 16.98 | 1.01 | 0.98 | 0.06 | 1.00 |
| **Year** | 1.27 | 1.14 | 1.53 | 1.13 | 0.79 | 0.65 | 0.88 |

**Table S24:** Collinearity assessment of the model investigating the factors associated with Leucocytozoon infection using urbanization level averaged per site | VIF = Variance Inflation Factor, CI = Confidence interval, SE = Standard Error. This assessment is based on the 'check_collinarity' function of the performance package.

| **Term** | **VIF** | **VIF_CI_low** | **VIF_CI_high** | **SE_factor** | **Tolerance** | **Tolerance**  **CI_low** | **Tolerance**  **CI_high** |
| --- | --- | --- | --- | --- | --- | --- | --- |
| **Site-level naturalness** | 1.17 | 1.07 | 1.43 | 1.08 | 0.85 | 0.70 | 0.94 |
| **Age** | 1.05 | 1.00 | 1.61 | 1.03 | 0.95 | 0.62 | 1.00 |
| **Sex** | 1.03 | 1.00 | 2.75 | 1.02 | 0.97 | 0.36 | 1.00 |
| **Year** | 1.16 | 1.06 | 1.42 | 1.08 | 0.86 | 0.71 | 0.94 |

###### 5.2.2.2. Distribution of residuals

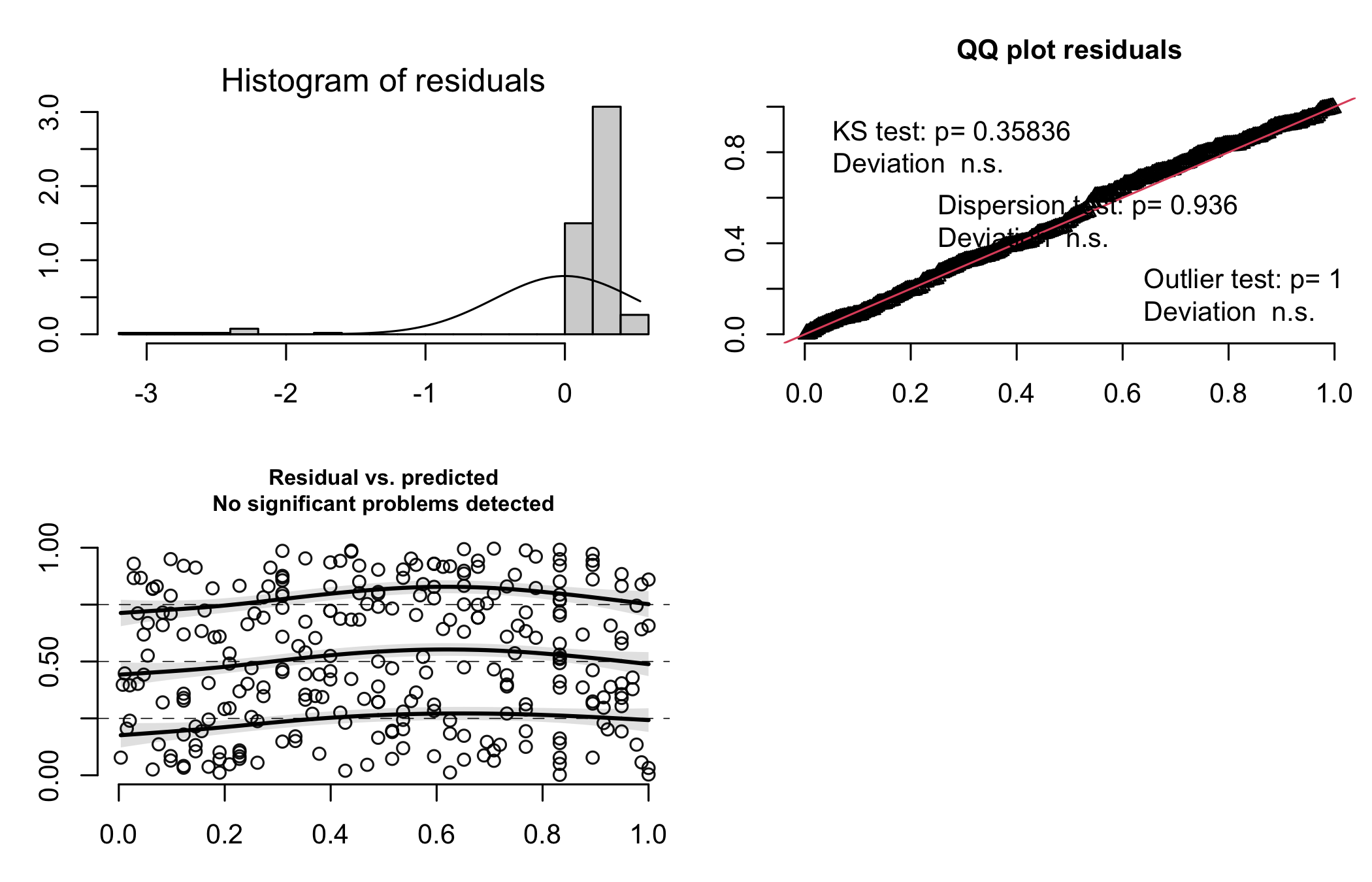

**Figure S21:** Model assumption check of the model on Plasmodium/Haemoproteus infection probability with urbanization level averaged per site | Depicted are the histogram of residuals, the Q-Q plot, and the scatter plot of the fitted values vs the residuals.

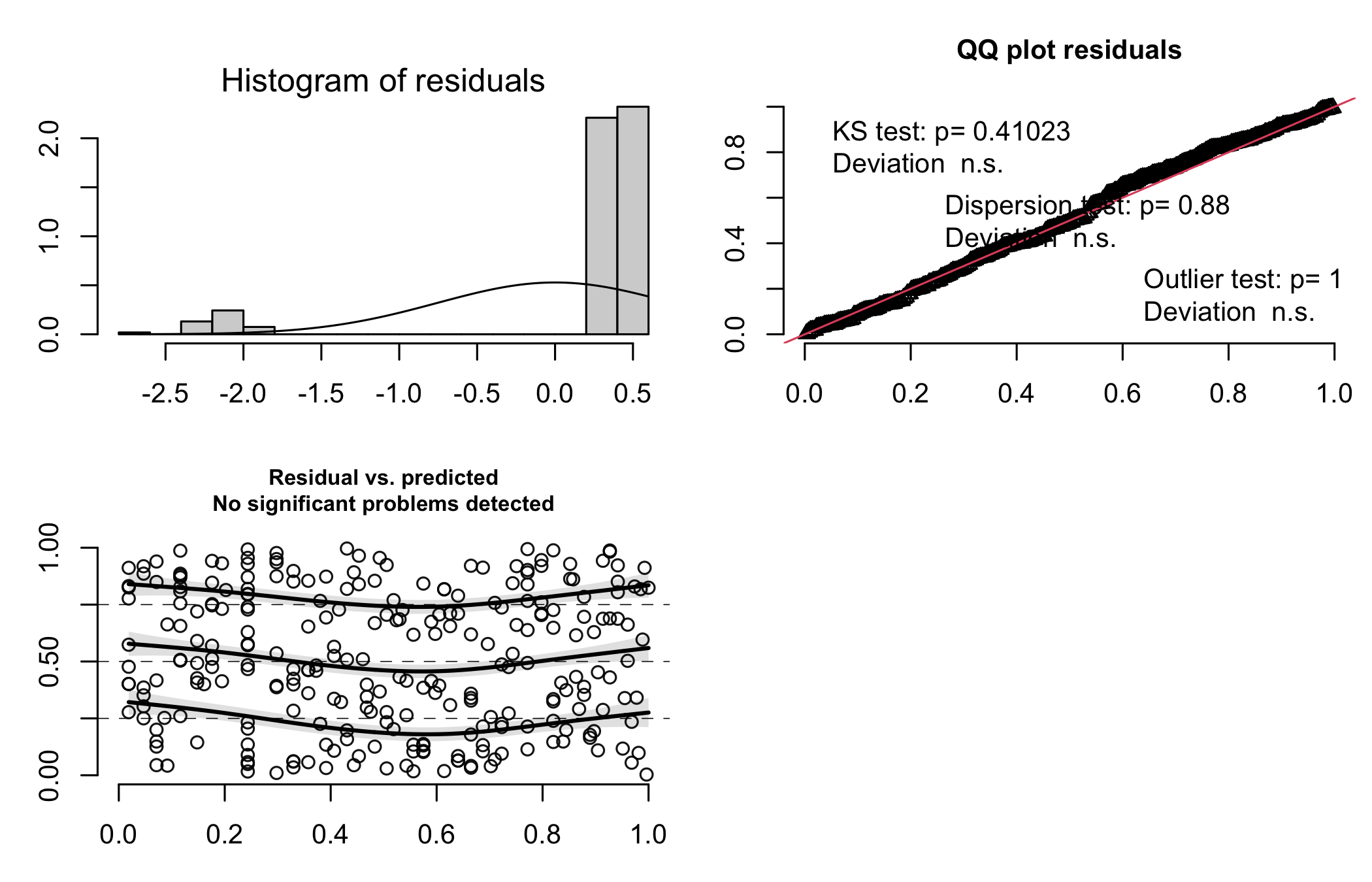

**Figure S22:** Model assumption check of the model on Leucocytozoon infection probability with urbanization level averaged per site | Depicted are the histogram of residuals, the Q-Q plot, and the scatter plot of the fitted values vs the residuals.

##### 5.2.3. Linear model stability

**Table S25:** Dfbetas of the model with urbanization level averaged per site on Plasmodium/Haemoproteus infection probability

| **Estimate** | **Min** | **Max** |
| --- | --- | --- |
| 4.11 | 3.93 | 4.94 |
| -0.21 | -0.37 | -0.12 |
| -0.31 | -0.75 | -0.22 |
| 0.73 | 0.57 | 0.86 |
| -0.94 | -1.13 | -0.77 |

**Table S26:** Dfbetas of the model with urbanization level averaged per site on Leucocytozoon infection probability.

| **Estimate** | **Min** | **Max** |
| --- | --- | --- |
| 1.47 | 1.27 | 1.74 |
| -0.02 | -0.06 | 0.06 |
| 0.21 | 0.04 | 0.27 |
| 0.26 | 0.18 | 0.33 |
| 0.84 | 0.72 | 0.95 |

#### 5.3. Urbanization as a gradient

##### 5.3.1. Linear model results

**Table S27:** Detailed output of fixed effects included in the GLM investigating *Plasmodium/Haemoproteus* prevalence in adults with urbanization level per nest-box.

|  | **Infection (Plasmodium/Haemoproteus)** | | | |
| --- | --- | --- | --- | --- |
| *Predictors* | *Log-Odds* | *std. Error* | *CI* | *p* |
| (Intercept) | 4.42 | 0.96 | 2.54 – 6.30 | - |
| Nest-level naturalness | -0.62 | 0.40 | -1.39 – 0.16 | 0.087 |
| Age | -0.30 | 0.34 | -0.96 – 0.36 | 0.395 |
| Sex [Male] | 0.69 | 0.74 | -0.75 – 2.14 | 0.336 |
| Year [2019] | -1.27 | 0.81 | -2.86 – 0.31 | 0.108 |
| **Random Effects** | | | | |
| σ^2^ | 3.29 | | | |
| τ_00_ _site_ | 0.00 | | | |
| N _site_ | 11 | | | |
| Observations | 267 | | | |

**Table S28:** Detailed output of fixed effects included in the GLM investigating *Leucocytozoon* prevalence in adults with urbanization level per nest-box.

|  | **Infection (Leucocytozoon)** | | | |
| --- | --- | --- | --- | --- |
| *Predictors* | *Log-Odds* | *std. Error* | *CI* | *p* |
| (Intercept) | -0.06 | 1.18 | -2.36 – 2.24 | - |
| Nest-level naturalness | 0.02 | 0.27 | -0.52 – 0.56 | 0.938 |
| Age | 0.24 | 0.34 | -0.43 – 0.91 | 0.471 |
| Sex [Male] | 0.54 | 0.52 | -0.48 – 1.55 | 0.293 |
| Year [2019] | 4.08 | 1.25 | 1.63 – 6.53 | **9.6x10^-6^** |
| **Random Effects** | | | | |
| σ^2^ | 3.29 | | | |
| τ_00_ _site_ | 4.70 | | | |
| ICC | 0.59 | | | |
| N _site_ | 11 | | | |
| Observations | 267 | | | |

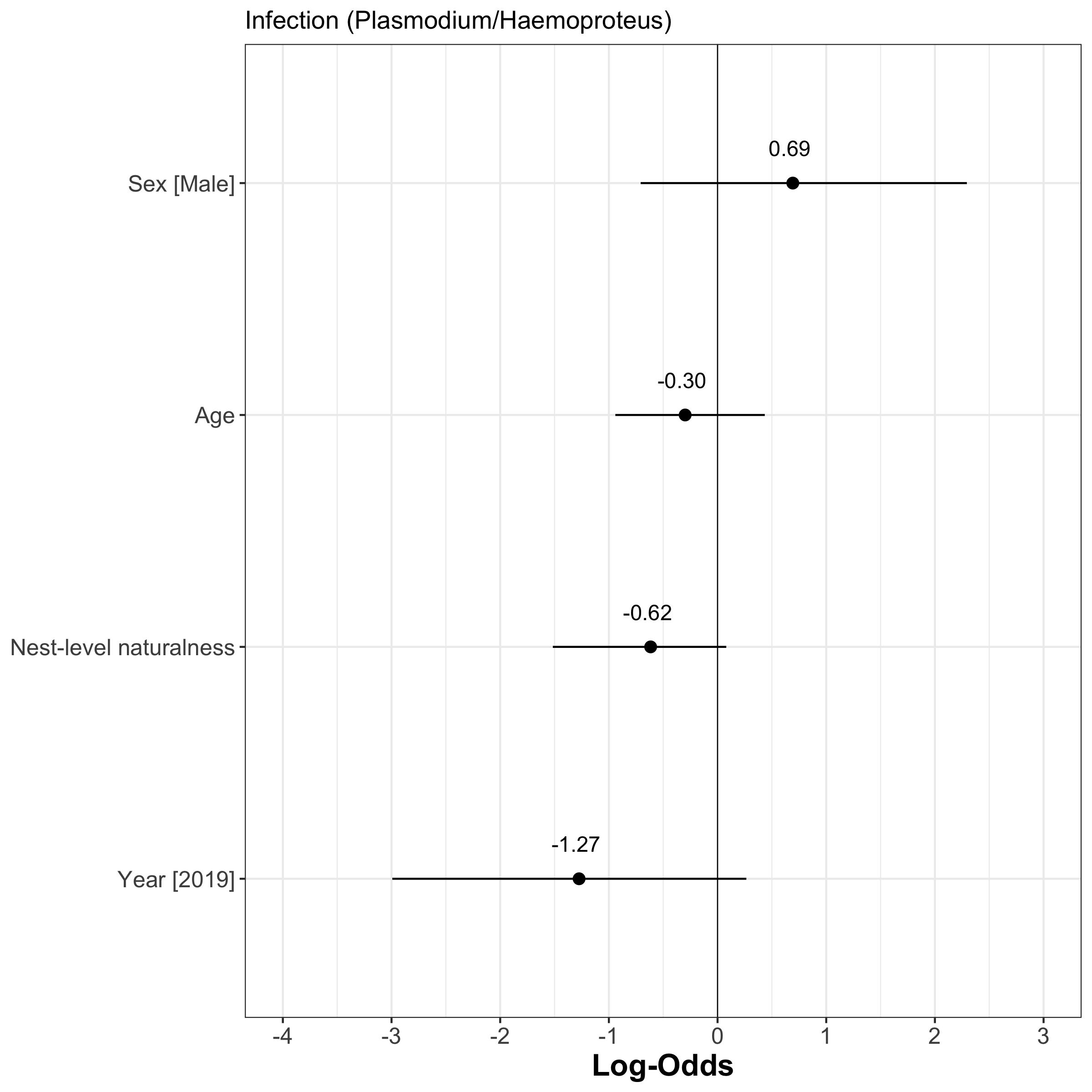

**Figure S22:** Forest plot of the estimates (Log-odds) of the model investigating the factors associated with Plasmodium/Haemoproteus infection using urbanization gradient at the nest-box level | The estimate value is indicated by plain points and its associate 95% confidence intervals by plain segments.

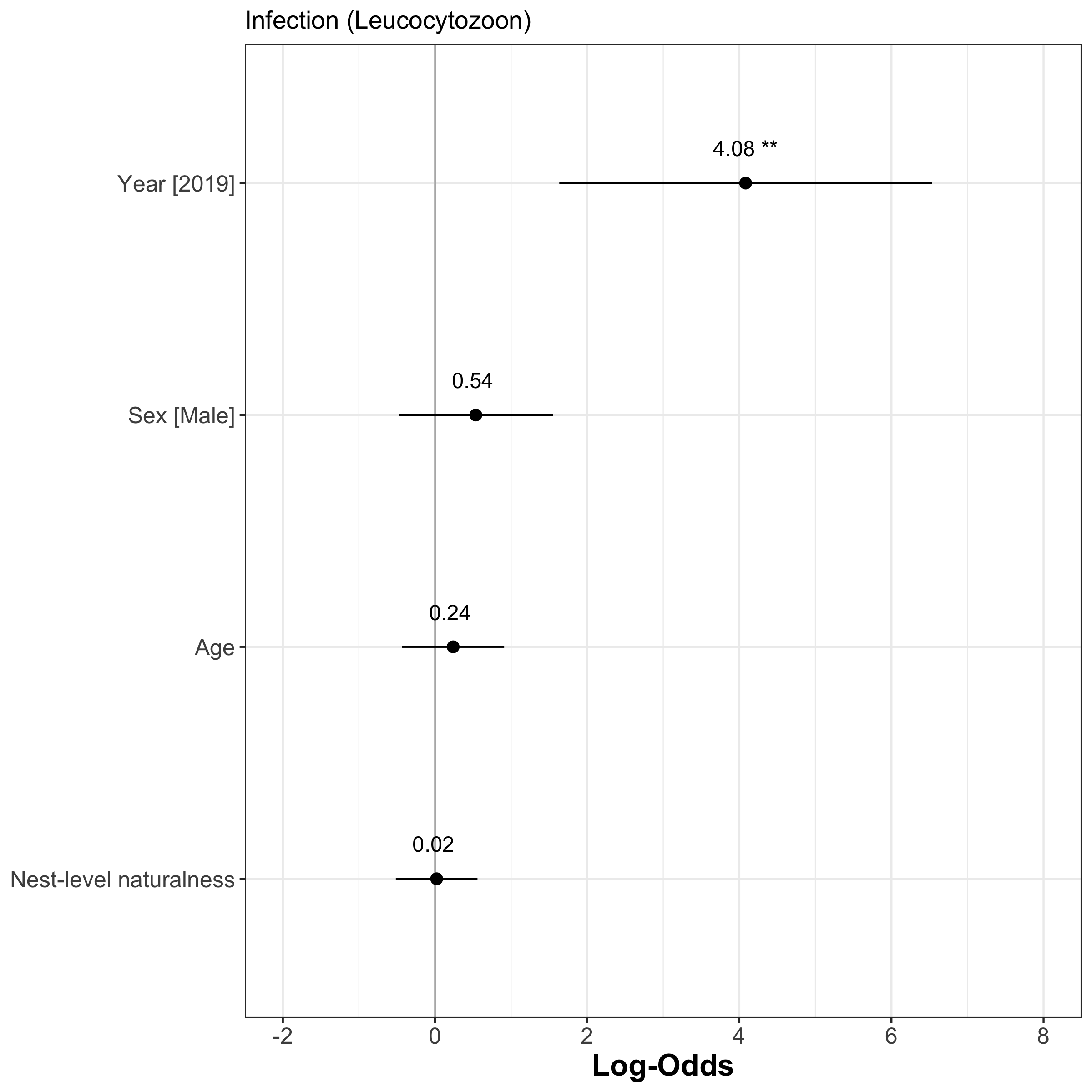

**Figure S23:** Forest plot of the estimates (Log-odds) of the model investigating the factors associated with Leucocytozoon infection using urbanization gradient at the nest-box level | The estimate value is indicated by plain points and its associate 95% confidence intervals by plain segments.

##### 5.3.2. Linear model assumptions

###### 5.3.2.1. Collinearity

**Table S29:** Collinearity assessment of the model investigating the factors associated with Plasmodium/Haemoproteus infection using the urbanization gradient at the nest-box level | VIF = Variance Inflation Factor, CI = Confidence interval, SE = Standard Error. This assessment is based on the 'check_collinarity' function of the performance package.

| **Term** | **VIF** | **VIF_CI_low** | **VIF_CI_high** | **SE_factor** | **Tolerance** | **Tolerance**  **CI_low** | **Tolerance**  **CI_high** |
| --- | --- | --- | --- | --- | --- | --- | --- |
| **Nest-level naturalness** | 1.19 | 1.08 | 1.45 | 1.09 | 0.84 | 0.69 | 0.92 |
| **Age** | 1.08 | 1.02 | 1.43 | 1.04 | 0.92 | 0.70 | 0.98 |
| **Sex** | 1.03 | 1.00 | 2.41 | 1.02 | 0.97 | 0.42 | 1.00 |
| **Year** | 1.25 | 1.12 | 1.50 | 1.12 | 0.80 | 0.67 | 0.89 |

**Table S30:** Collinearity assessment of the model investigating the factors associated with Leucocytozoon infection using the urbanization gradient at the nest-box level| VIF = Variance Inflation Factor, CI = Confidence interval, SE = Standard Error. This assessment is based on the 'check_collinarity' function of the performance package.

| **Term** | **VIF** | **VIF_CI_low** | **VIF_CI_high** | **SE_factor** | **Tolerance** | **Tolerance**  **CI_low** | **Tolerance**  **CI_high** |
| --- | --- | --- | --- | --- | --- | --- | --- |
| **Nest-level naturalness** | 1.01 | 1.00 | 1823.57 | 1.00 | 0.99 | 0.00 | 1.00 |
| **Age** | 1.03 | 1.00 | 3.59 | 1.01 | 0.97 | 0.28 | 1.00 |
| **Sex** | 1.04 | 1.00 | 1.98 | 1.02 | 0.96 | 0.51 | 1.00 |
| **Year** | 1.02 | 1.00 | 4.46 | 1.01 | 0.98 | 0.22 | 1.00 |

###### 5.3.2.2. Distribution of residuals

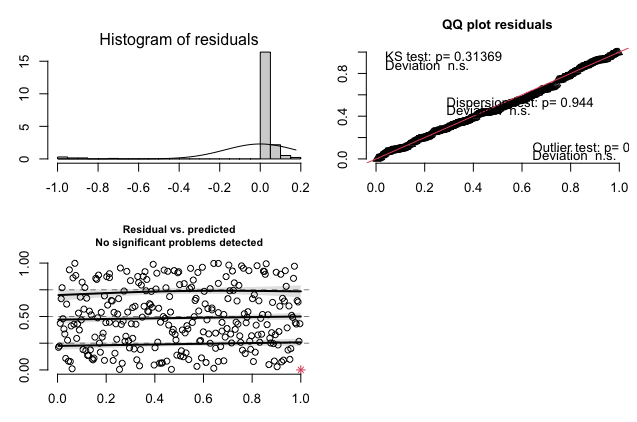

**Figure S24:** Model assumption check of the model on Plasmodium/Haemoproteus infection probability with the urbanization gradient at the nest-box level | Depicted are the histogram of residuals, the Q-Q plot, and the scatter plot of the fitted values vs the residuals.

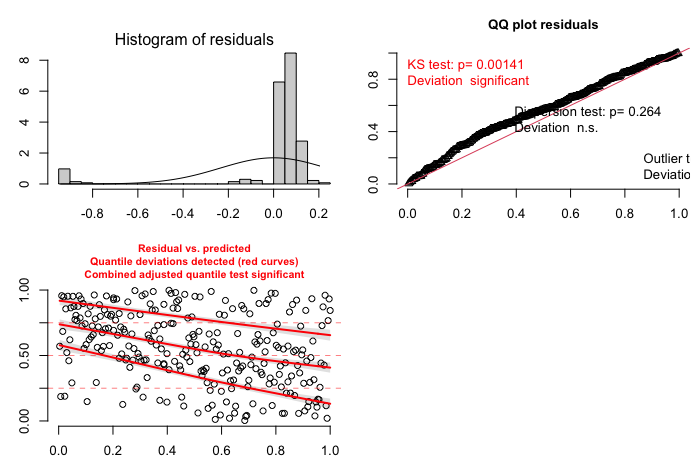

**Figure S25:** Model assumption check of the model on Leucocytozoon infection probability with urbanization gradient at the nest-box level the urbanization gradient at the nest-box level | Depicted are the histogram of residuals, the Q-Q plot, and the scatter plot of the fitted values vs the residuals.

##### 5.3.3. Linear model stability

**Table S31:** Dfbetas of the model with the urbanization gradient at the nest-box level on Plasmodium/Haemoproteus infection probability

| **Estimate** | **Min** | **Max** |
| --- | --- | --- |
| 4.42 | 3.46 | 5.37 |
| -0.62 | -1.07 | 0.70 |
| -0.30 | -1.45 | 0.17 |
| 0.69 | 0.12 | 1.01 |
| -1.27 | -1.74 | -0.25 |

**Table S32:** Dfbetas of the model with the urbanization gradient at the nest-box level on Leucocytozoon infection probability

| **Estimate** | **Min** | **Max** |
| --- | --- | --- |
| -0.6 | -0.33 | 0.77 |
| 0.02 | -0.36 | 0.67 |
| 0.24 | -1.09 | 0.56 |
| 0.54 | 0.25 | 0.77 |
| 4.08 | 3.60 | 4.30 |

### 6. Packages information

**Table S33:** R packages used for analysis | For packages directly available in R (the core of base packages), no version is associated as it depends on the R version used. For those analyses, the R version used was v4.2.1

| **Package** | **Version** | **Reference** |
| --- | --- | --- |
| ape | 5.6-2 | Paradis E, Schliep K (2019). "ape 5.0: an environment for modern phylogenetics and evolutionary analyses in R." *Bioinformatics*, *35*, 526-528. |
| base | - | R Core Team (2022). *R: A Language and Environment for Statistical Computing*. R Foundation for Statistical Computing, Vienna, Austria. https://www.R-project.org/. |
| BiodiversityR | 2.14-4 | Kindt R, Coe R (2005). *Tree diversity analysis. A manual and software for common statistical methods for ecological and biodiversity studies*. World Agroforestry Centre (ICRAF), Nairobi (Kenya). ISBN 92-9059-179-X, http://www.worldagroforestry.org/output/tree-diversity-analysis. |
| datasets | - | R Core Team (2022). *R: A Language and Environment for Statistical Computing*. R Foundation for Statistical Computing, Vienna, Austria. https://www.R-project.org/. |
| DHARMa | 0.4.6 | Hartig F (2022). *DHARMa: Residual Diagnostics for Hierarchical (Multi-Level / Mixed) Regression Models*. R package version 0.4.6, https://CRAN.R-project.org/package=DHARMa. |
| dplyr | 1.0.10 | Wickham H, François R, Henry L, Müller K (2022). dplyr: A Grammar of Data Manipulation. R package version 1.0.10, https://CRAN.R-project.org/package=dplyr. |
| ggplot2 | 3.4.1 | Wickham H (2016). *ggplot2: Elegant Graphics for Data Analysis*. Springer-Verlag New York. ISBN 978-3-319-24277-4, https://ggplot2.tidyverse.org. |
| graphics | - | R Core Team (2022). *R: A Language and Environment for Statistical Computing*. R Foundation for Statistical Computing, Vienna, Austria. https://www.R-project.org/. |
| grDevices | - | R Core Team (2022). *R: A Language and Environment for Statistical Computing.* R Foundation for Statistical Computing, Vienna, Austria. https://www.R-project.org/. |
| huxtable | 5.5.2 | Hugh-Jones D (2022). *huxtable: Easily Create and Style Tables for LaTeX, HTML and Other Formats*. R package version 5.5.2, https://CRAN.R-project.org/package=huxtable. |
| lattice | 0.20-45 | Sarkar D (2008). *Lattice: Multivariate Data Visualization with R*. Springer, New York. ISBN 978-0-387-75968-5, http://lmdvr.r-forge.r-project.org. |
| lme4 | 1.1-30 | Bates D, Mächler M, Bolker B, Walker S (2015). Fitting Linear Mixed-Effects Models Using lme4. *Journal of Statistical Software*, *67*(1), 1-48. doi:10.18637/jss.v067.i01 https://doi.org/10.18637/jss.v067.i01. |
| lmerTest | 3.1-3 | Kuznetsova A, Brockhoff PB, Christensen RHB (2017). lmerTest Package: Tests in Linear Mixed Effects Models. *Journal of Statistical Software*, *82*(13), 1-26. doi:10.18637/jss.v082.i13 https://doi.org/10.18637/jss.v082.i13. |
| lmtest | 0.9-40 | Zeileis A, Hothorn T (2002). Diagnostic Checking in Regression Relationships. *R News*, *2*(3), 7-10. https://CRAN.R-project.org/doc/Rnews/. |
| Matrix | 1.5-3 | Bates D, Maechler M, Jagan M (2022). Matrix: Sparse and Dense Matrix Classes and Methods. R package version 1.5-3, https://CRAN.R-project.org/package=Matrix. |
| methods | - | R Core Team (2022). *R: A Language and Environment for Statistical Computing*. R Foundation for Statistical Computing, Vienna, Austria. https://www.R-project.org/. |
| performance | 0.10.2 | Lüdecke D, Ben-Shachar M, Patil I, Waggoner P, Makowski D (2021). performance: An R Package for Assessment, Comparison and Testing of Statistical Models. *Journal of Open Source Software*, *6*(60), 3139. doi:10.21105/joss.03139 https://doi.org/10.21105/joss.03139. |
| permute | 0.9-7 | Simpson G (2022). *permute: Functions for Generating Restricted Permutations of Data*. R package version 0.9-7, https://CRAN.R-project.org/package=permute. |
| reshape | 0.8.9 | Wickham H (2007). Reshaping data with the reshape package. *Journal of Statistical Software*, *21*(12). https://www.jstatsoft.org/v21/i12/. |
| sf | 1.0-9 | Pebesma E (2018). Simple Features for R: Standardized Support for Spatial Vector Data. *The R Journal*, *10*(1), 439-446. doi:10.32614/RJ-2018-009, https://doi.org/10.32614/RJ-2018-009. |
| sjPlot | 2.8.12 | Lüdecke D (2022). *sjPlot: Data Visualization for Statistics in Social Science*. R package version 2.8.12, https://CRAN.R-project.org/package=sjPlot. |
| stats | - | R Core Team (2022). *R: A Language and Environment for Statistical Computing*. R Foundation for Statistical Computing, Vienna, Austria. https://www.R-project.org/. |
| tcltk | - | R Core Team (2022). *R: A Language and Environment for Statistical Computing.* R Foundation for Statistical Computing, Vienna, Austria. https://www.R-project.org/. |
| tidyr | 1.2.1 | Wickham H, Girlich M (2022). t*idyr: Tidy Messy Data*. R package version 1.2.1, https://CRAN.R-project.org/package=tidyr. |
| utils | - | R Core Team (2022). *R: A Language and Environment for Statistical Computing*. R Foundation for Statistical Computing, Vienna, Austria. https://www.R-project.org/. |
| vegan | 2.6-2 | Oksanen J, Simpson G, Blanchet F, Kindt R, Legendre P, Minchin P, O'Hara R, Solymos P, Stevens M, Szoecs E, Wagner H, Barbour M, Bedward M, Bolker B, Borcard D, Carvalho G, Chirico M, De Caceres M, Durand S, Evangelista H, FitzJohn R, Friendly M, Furneaux B, Hannigan G, Hill M, Lahti L, McGlinn D, Ouellette M, Ribeiro Cunha E, Smith T, Stier A, Ter Braak C, Weedon J (2022). *vegan: Community Ecology Package*. R package version 2.6-2, https://CRAN.R-project.org/package=vegan. |
| zoo | 1.8-10 | Zeileis A, Grothendieck G (2005). zoo: S3 Infrastructure for Regular and Irregular Time Series. *Journal of Statistical Software*, *14*(6), 1-27. doi:10.18637/jss.v014.i06 https://doi.org/10.18637/jss.v014.i06. |
